# Integrated Closed-loop Control of Bio-actuation for Proprioceptive Bio-hybrid Robots

**DOI:** 10.1101/2024.01.30.577987

**Authors:** Miriam Filippi, Aiste Balciunaite, Antonia Georgopoulou, Pablo Paniagua, Felix Drescher, Minghao Nie, Shoji Takeuchi, Frank Clemens, Robert Katzschmann

## Abstract

Biohybrid robots are emergent soft robots that combine engineered artificial structures and living biosystems to exploit unique characteristics of biological cells and tissues. Skeletal muscle tissue-based bio-actuators can respond to externally applied stimuli, such as electrical fields. However, current bio-actuation systems rely on open-loop control strategies that lack knowledge of the actuator’s state. The regulation of output force and position of bio-hybrid robots requires self-sensing control systems that combine bio-actuators with sensors and control paradigms. Here, we propose a soft, fiber-shaped mechanical sensor based on a composite with piezoresistive properties that efficiently integrates with engineered skeletal muscle tissue and senses its contracting states in a cell culture environment in the presence of applied electrical fields. After testing the sensor’s insulation and biocompatibility, we characterized its sensitivity for typical strains (<1%) and proved its ability to detect motions from contractile skeletal muscle tissue constructs. Finally, we showed that the sensor response can feed an autonomous control system, thus demonstrating the first proprioceptive bio-hybrid robot that can sense and respond to its contraction state. In addition to inspiring intelligent implantable systems, informative biomedical models, and other bioelectronic systems, the proposed technology will encourage strategies to exceed the durability, design, and portability limitations of biohybrid robots and confer them decisional autonomy, thus driving the paradigm shift between bio-actuators and intelligent bio-hybrid robots.

**One Sentence Summary:** Integrating soft mechanical sensors into engineered skeletal muscle tissue enables bio-hybrid robots with proprioception.

## 1. Introduction

Bio-hybrid robots are dynamic machines that use the unique properties of living cells to achieve controlled motion (*1*). Biohybrid robotics mainly focuses on realizing bio–actuators engineered from cardiac or skeletal muscle cells (*1*, *2*). Muscle cells are combined with scaffolding materials to grow in different 3D geometries. The resulting bio-actuators are assembled with other synthetic, deformable components to form robots capable of various functions, including locomotion, gripping, and pumping. By exploiting the specialized contractility of skeletal or cardiac muscle cells to induce deformation forces, bio-actuators have the potential to overcome the inability of synthetic materials to replicate native muscle’s unique functions, such as mechanical compliance, flexibility, sensing capabilities, self-healing, and adaptability (*3*, *4*).

Even if various aspects concerning the scalability and dynamicity of muscle bio-actuators have been explored (*5–8*), the motion control paradigm only relies on an active input delivery, in which electrical fields or optical signals start or synchronize the contraction of natural or optogenetically modified cells, respectively (*9*, *10*). The contraction response depends on pre-defined stimulation paradigms, whose optimized parametrization is highly specific to the investigated biosystem. While pre-defined, open-loop stimulation of engineered muscle tissue benefits the initial development and understanding of bio-hybrid robots (*11*, *12*), it only allows for static bio-actuator control, hindering integration into intelligent, autonomous systems. Adaptive and intelligent bio-hybrid robots must incorporate closed-loop control by integrating bio-actuators and sensors capable of understanding their biomechanical state and subsequently regulating the actuation responses. Feedback control systems for bio-hybrid robotics do not yet exist.

In general, the development of tissue-integrated bioelectronics is primarily hindered by technical hurdles that reduce the electronics’ operativity and durability within bioengineered systems. A promising emerging sensor technology for tissue integration is piezoresistive mechanical sensors. Unfortunately, these materials display limited biocompatibility, reduced operativity in wet environments, and a significant mechanical mismatch with the soft target tissue. Other factors that limit the biomedical usability of these sensors are electrolysis of biological liquid media and conductive interference from the application of additional electrical currents, such as those typically used to stimulate bio-actuators. Manufacturing insulated piezoresistive mechano-sensors for bio-actuators from biocompatible and soft materials could overcome these challenges.

Here, we present a tissue-integrated piezoresistive sensor that due to its mechanical compliance, biocompatibility, and sensitivity can be integrated into developing bio-actuators and detect their stimulated motion responses. Our sensor was fabricated via thermoplastic extrusion from an efficiently manufacturable and biocompatible composition of carbon black (CB) and medical-grade styrene-based copolymers. To apply the sensor to biomedical systems, we designed it as a fiber with a small diameter (100 µm) and high softness (20A shore hardness). Furthermore, we electrically insulated the fiber with a biocompatible and bioadhesive coating to operate effectively within electrolytic media. We characterized the sensor functionality via mechanoelectrical characterization and proved its biocompatibility and integration with skeletal muscle tissue-based bio-actuators. We showed that our integrated sensor detects bio-actuator motions with tens-of-micrometers lateral displacements and forces in the few-mN ranges, which are performance parameters expected for state-of-the-art bio-actuators.(*1*, *9*, *13*) Finally, we demonstrated the first proprioceptive bio-hybrid robot capable of understanding and controlling its contraction state by applying a feedback control system on data from a sensor fiber integrated into our bio-actuator.

## 2. Results

### 2.1 Sensorizing the Engineered Skeletal Muscle

To sensorize our engineered skeletal muscle tissue, we developed a thin, stretchable, fiber-shaped sensor based on a piezoresistive CB-polymer composite that was coated with a bioadhesive insulator layer (**Fig. 1A**). We characterized the mechano-electrical profile of the sensor, and its sensing performance in biologically-relevant environments and when embedded in soft matrices and engineered skeletal muscle tissue. The sensor was used to measure strain-type mechanical loading deriving from the contraction response of electrically stimulated skeletal muscle-based bio-actuators. We realized a threshold-based control mechanism to stop the bio-actuator contraction autonomously, thus showcasing a bio-hybrid robot capable of autonomously sensing and controlling its contraction state (**Fig. 1B**).

**Figure 1.**
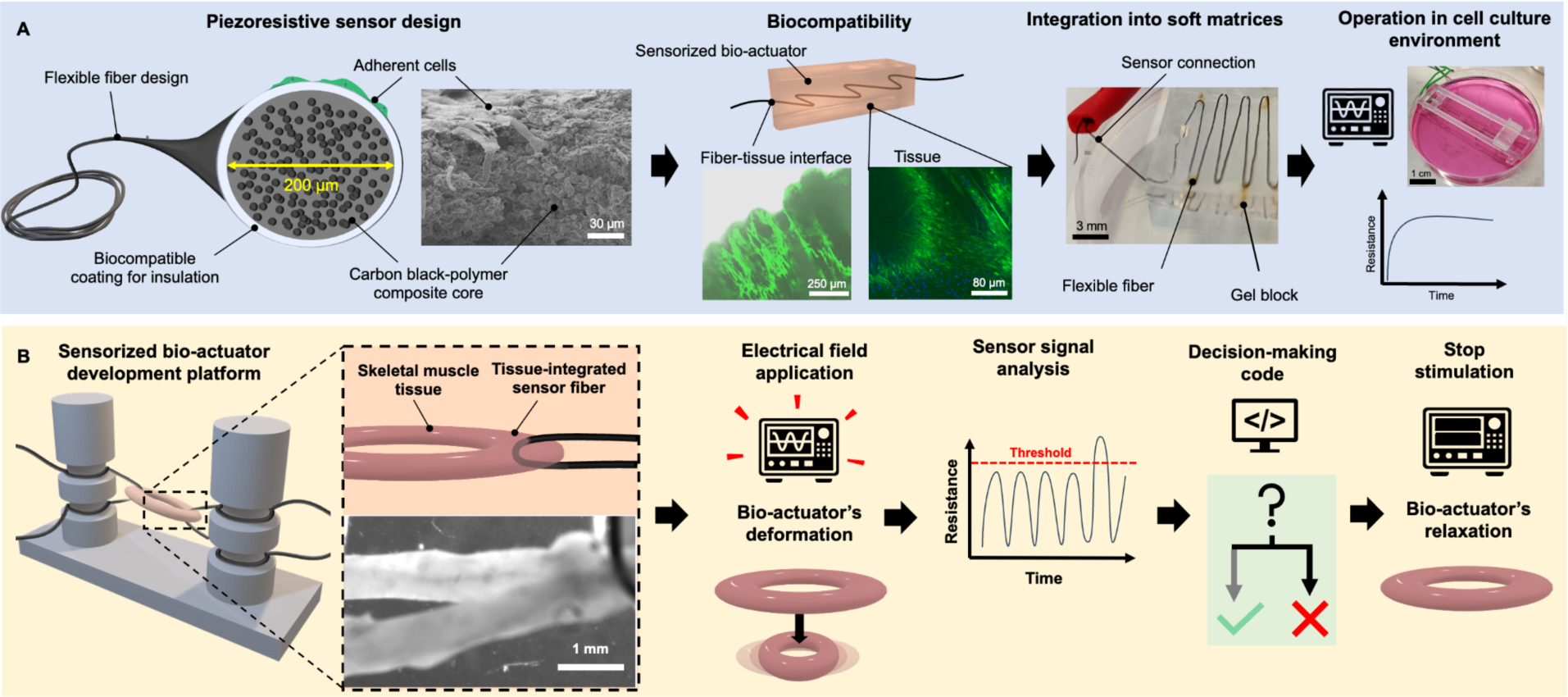
Realization of a proprioceptive system in a skeletal muscle tissue-based bio-actuator. (**A**) The mechano-sensor uses a piezoresistive composite designed as a styrene-insulated fiber. The fiber was characterized by its sensing performance, integrability in soft matrices, biocompatibility in engineered skeletal muscle tissue, and operativity in a cell culture environment. (**B**) When assembled with bio-actuators, the sensor revealed micromotions triggered by electrical stimulation without disturbance from applied electrical fields, and the sensor signal could be used to modulate the stimulation pulses provided by muscle electrical stimulation setup.

### 2.2 Manufacturing of the sensor and its properties

We produced the piezoresistive fiber of the sensor with thermoplastic processing techniques that involved high-shear mixing and melt extrusion of a composite based on carbon black and implant-grade styrene-based copolymer (medTPS) (**Fig. 2A**). To maximize the mechanical affinity with the target soft tissue, reduce strain shielding effects, and guarantee an efficient strain transfer from the tissue to the fiber (*14*, *15*), we aimed to fabricate our sensor with low Shore hardness materials, which can coherently integrate within soft matter (such as body tissue, engineered tissue, and 3D cell culture models) (*16–18*). We investigated melt-extruded fibers with different Shore hardness medTPS (20, 40, and 50A) to define the lowest Shore hardness values of medTPS that would enable the successful fabrication of the system.

**Figure 2.**
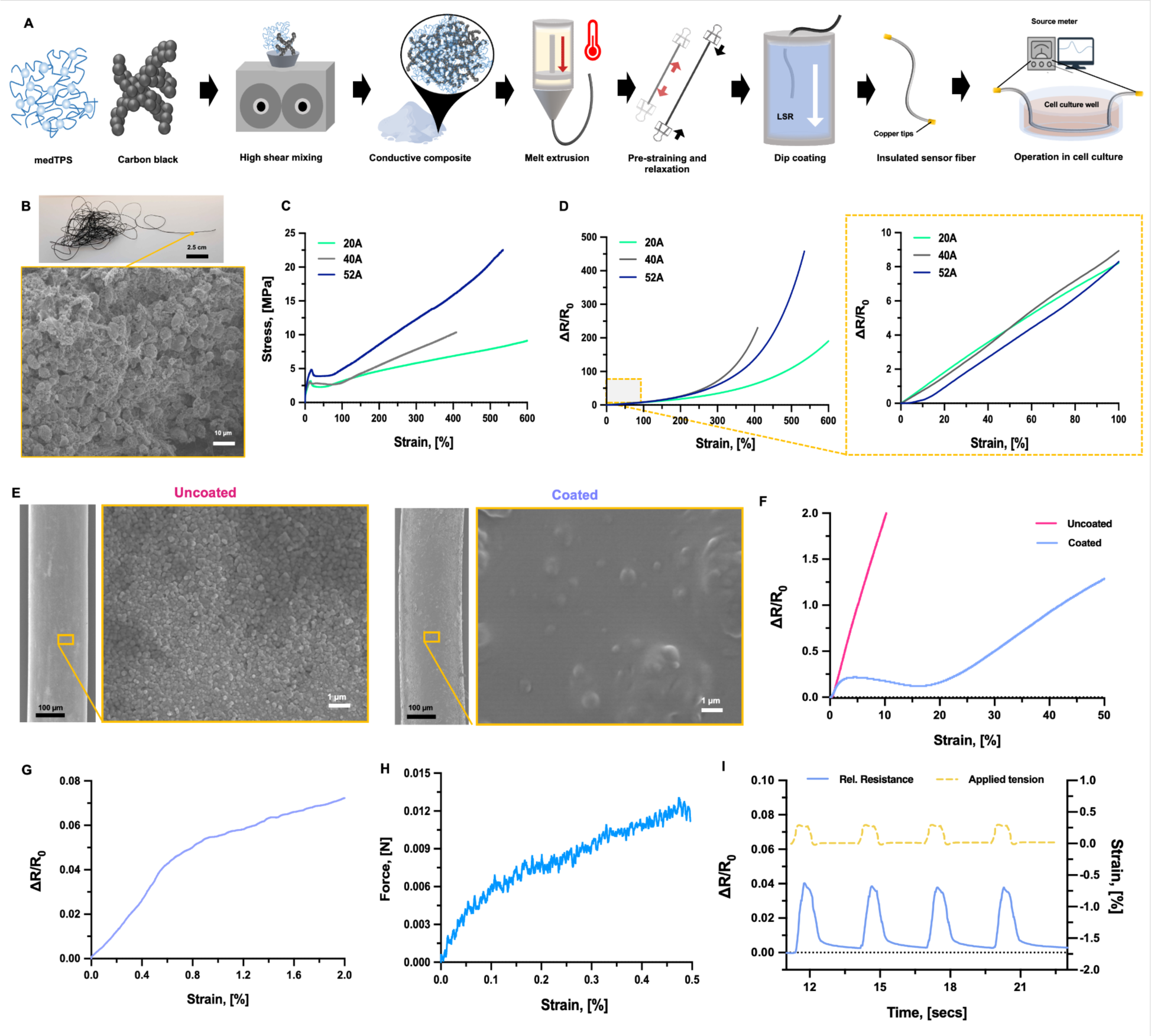
Manufacturing and performance of the sensor. **(A)** Thermoplastic sensor fabrication from carbon black and styrene-based copolymers that were high-shear mixed and heated extruded to form a fiber-shaped system. The fiber was pre-strained and then coated to remain functional within cell culture and stimulation environments. (**B**) Optical picture of the sensor fiber and SEM of its internal composite structure. Strain-dependent variation of the stress (**C**) and the relative resistance (ΔR/R_0_) (**D**) of sensors composed of polymers with different Shore hardness values (20, 40, and 52 A). The dashed orange line indicates the magnified inset of ΔR/R_0_ variation for the 0-100% strain range. (**E**) Scanning Electron Microscopy pictures of the uncoated and coated fibers. Variation of the relative resistance (ΔR/R_0_) of coated and uncoated sensors dependent on the strain for large (up to 50%) (**F**) and small (0-2%) strain values (**G**). (**H**) Force over the applied strain for small strain values. (**I**) Sensor dynamic response: time-dependent variation of the relative resistance of the coated fibers (blue line) under repeated strain stress applied at 0.2%.

We manufactured the fibers by extruding their component materials through a 0.2 mm diameter nozzle. The resulting fiber had an internal composite appearance (**Fig. 2B**) and was then characterized with mechano-electrical tests to the point of fracture (**Fig. 2C**). The sensor fiber with the lowest Shore hardness (20A) reached the highest elongation at the point of fracture and displayed the lowest ultimate strength. All the fiber compositions displayed necking at the yield point, which is typical of TPS composites with ultra-high CB concentration (50 w/w %) (*19*, *20*). The yield point appeared at 24% strain and was not affected by the Shore hardness of the medTPS.

As for the response of the electrical resistance, the sensor fibers exhibited a positive piezoresistive response (**Fig. 2D**). The resistance increased sharply at high strains (>300%). In contrast with other fibers, the fiber with the highest Shore hardness (50A) showed almost no change in the relative resistance for low strains (below 10%). At 10% strain, the response of the 20A and 40A fibers displayed a positive slope with a Gauge Factor (GF) of 10. At 50 % strain, the GF for the 50A fiber and the other two fiber types was 6 and 8.4, respectively. As the 20A fiber displayed high stretchability and sensitivity at low strains, this was selected for all further experiments.

After fabrication, we pre-strained the fiber to make it thinner and then we used a dip coating method with liquid silicone rubber to form an electrical insulation layer around it. Scanning electron microscopy (SEM) revealed that the pre-strained fibers had a diameter close to the extrusion nozzles size (220.5± 22 µm), and the dip coating process resulted in a 4-μm-thick layer (**Fig. 2E**). The relative resistance variation of the coated 20A fibers (**Fig. 2F** and **S1-2**) was monotonic for strain stresses larger than 20% and comprised in the 0-5% range.

To understand if our sensor could detect minimal displacements likely caused by small output forces of bio-actuators (*9*, *13*, *21*), we investigated the sensor behavior in the 0-2% strain ranges, finding that the stress and relative resistance response coherently reacted to the applied strains (**Fig. 2G** and **S3-7**). With 0-3% strain deformation, the coated fiber provided a monotonic response (relative resistance range: 0-0.003) (**Fig. 2G-H**). The force detection was in the range expected for bio-actuators (few mN range) (*9*, *21*). The same behavior emerged in a dynamic test that mimicked a typical bio-actuator’s stimulation paradigm (0.2% strain stress applied at 1Hz in **Fig. 2I**). Thus, we expected the sensor to operate consistently during the mechanostimulation of a bio-actuator. Moreover, pre-straining the fiber before coating augmented the GF as calculated under small (1-2%) strains application (**Fig. S8**), thus suggesting that using pre-strained fibers could serve to generate sensitive sensors that are useful to operate on bio-actuators.

### 2.3 Sensor biocompatibility with muscle cell culture and tissue formation

To understand the applicability of our sensor fiber as a bioelectronic material for cell culture models and tissue-embedded applications, we tested its biocompatibility. First, by a Resazurin assay, we assessed the effects of the fiber constituent materials on the viability of murine myoblasts, grown as a monolayer and then exposed to different materials concentrations (0.1-100 mg/mL) and for variable time ranges (up to 14 days) (**Fig. 3A**). At any tested condition, cell viability was above 85%, demonstrating that the fiber is highly biocompatible in both its coated and uncoated designs (**Fig. 3B** and **S9**), and preserves cell proliferation ability (**Fig. S10**). As observed with Live/Dead assays and Scanning Electron Microscopy, cells could adhere to the coated sensor’s surface (**Fig. 3C-D**). After five days of growth, the cells formed a biofilm on the fiber surface. After ten days, the biofilm covered the fiber almost in its entirety and contained cells stacked into multiple layers and only sporadic apoptotic bodies (**Fig. 3D**). In addition to skeletal muscle cells, other cell types could adhere and grow on the fibers (**Fig. S12**), suggesting that the sensor has the potential to form effective bio-interfaces with various cells and be used for different bioelectronic systems.

**Figure 3.**
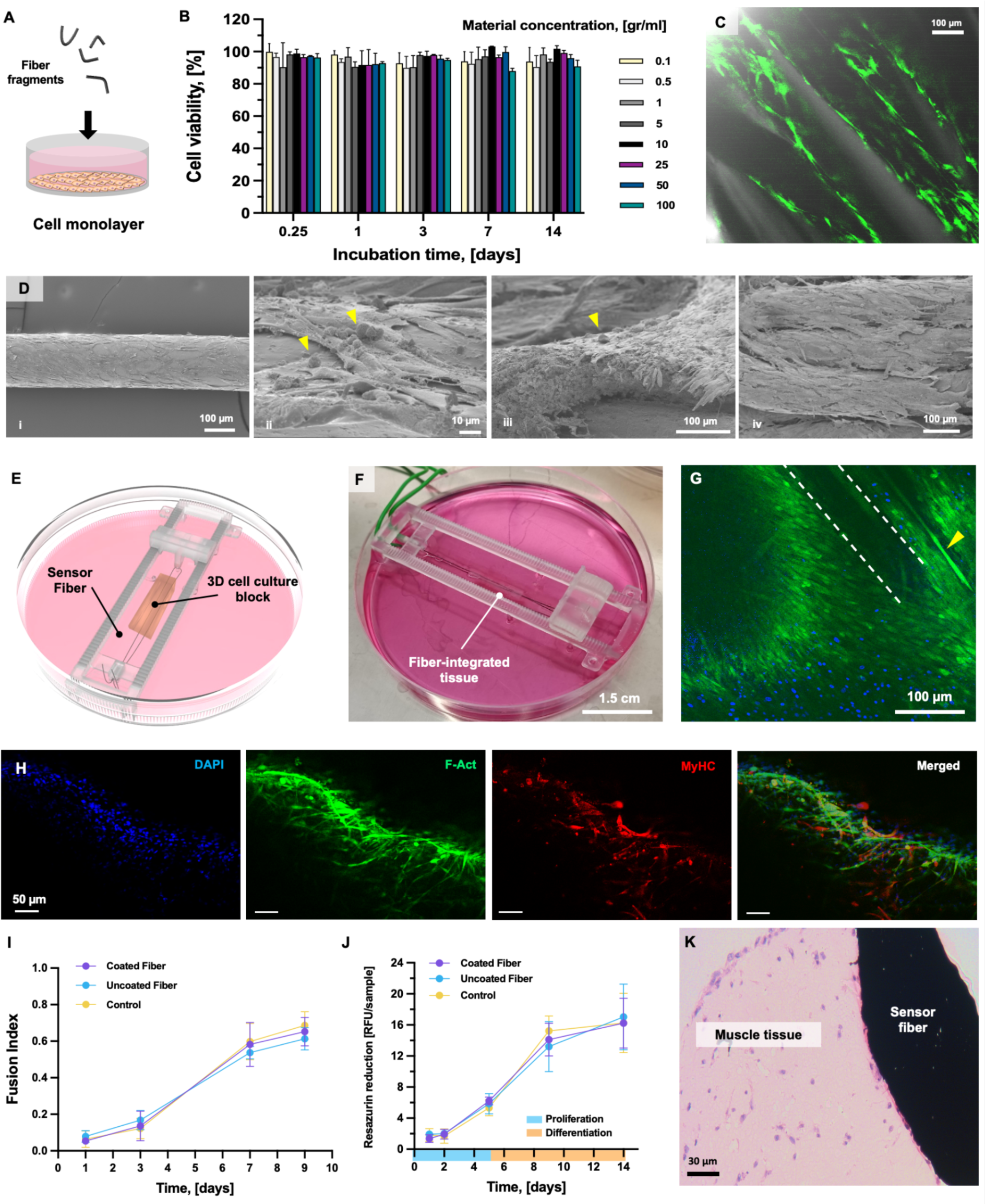
Biocompatibility of the tissue-integrated fiber. **(A)** Scheme of the incubation assay of a myoblast cell monolayer with sensor fragments. **(B)** Cell viability was measured via a Resazurin assay after exposure to different concentrations of the constituent materials of the coated fibers. **(C)** Confocal imaging of Live/Dead staining of murine myoblasts adhering to the fiber surface**. (D)** Scanning Electron Microscopy of the coated fiber seeded with myoblasts on days 5 (i, ii, iii) and 10 (iv). Bodies with rounded morphologies (apoptotic bodies) are shown with yellow arrows. (**E**) Schematical setup for functional tests of the sensor integrated with a cm-scale 3D cell culture model. (**F**) Actual test setup for the sensor integrated with a cm-scale 3D cell culture model. (**G**) Confocal imaging of a construct stained with Live/Dead assay, revealing viable myotubes (yellow arrow) aligned to the fiber’s orientation (dashed lines). (**H**) Immunofluorescence staining of myogenic differentiation markers on the skeletal muscle tissue (day 15). Nuclei, F-actin, and MyHC, are shown in blue, green, and red, respectively. **(I)** Cell metabolic rate over time measured from Resazurin assay from constructs with a coated, uncoated, or no fiber embedded. (**J**) Fusion index (*i.e*., number of nuclei inside MyHC-positive myotubes divided by the total number of nuclei present in a field of view) calculated from confocal imaging (n = 3). (**K**) H&E staining of the skeletal muscle tissue reveals cells dispersed in the matrix and a coherent interface developed with the integrated fiber.

To understand if the sensor could be integrated with a developing 3D tissue model, we embedded the coated fiber into a myoblast-laden hydrogel construct (1×0.5×0.5 cm) during the biofabrication process (**Fig. 3E-F** and **S12-14**). We analyzed the cell proliferation, differentiation ability, and the skeletal muscle tissue maturation process. When observing the coated fiber-embedded tissue in confocal microscopy, high-density myotubes with parallel orientation to the fiber direction were found (**Fig. 3G**), which stained positive for the late differentiation marker Myosin Heavy Chain (MyHC) (**Fig. 3H**), featuring a length (around 180 µm) and fusion index (0.6 ca.) that were not statistically different from the value ranges observed in the control constructs (245 µm and 0.6, respectively) (**Fig. 3I** and **S15**). The cells in the fiber-embedded tissue constructs followed the same proliferation trend observed for cells in control tissue constructs (**Fig. 3J**), with a fast proliferation phase in the initial culture time (first week) and a slow proliferation kinetics expected for the differentiation phase of the culture protocol (from day 8 to 14). The secretome analysis revealed that the production of myogenic function regulators was not affected by the presence of the fiber (**Fig. S15**). The living tissue is tightly attached to the fiber, forming a compact biphasic system with the potential to efficiently transfer a mechanical load for optimized sensing (**Fig. 3K**). The interface between fiber and matrix in the absence of cells was loose, suggesting that the co-developed living tissue plays an active role in stabilizing the interface (**Fig. S16-18**).

### 2.4 Operation of the sensor in biologically relevant environments

We aimed to understand how the sensor would operate when exposed to biologically relevant environments. Therefore, we tested the sensor’s reliability when integrated into soft matrices mimicking soft tissues. We also cultured long-term in cell culture conditions and media. We also tested the sensor embedded in tissue in the presence of applied electrical fields.

To understand how embedding the fiber in a soft matrix affected the sensor response under realistic loading conditions (tension, compression), we integrated the sensor in soft tissue-mimicking phantoms (**Fig. 4A-C, S19-20**). If embedded in a straight configuration (**Fig. 4A**), the sensor did not reliably respond to tensile stress, likely due to an inefficiently stabilized interface between the matrix and the sensor, which resulted in the fiber slipping within the non-cellularized phantoms. Therefore, we mechanically stabilized the sensor integration by arranging the fiber in a pre-tensioned serpentine design that could remain embedded in the gel block during applied mechanical stress (**Fig. 4C**). Even if in both configurations, the fiber compliantly bent with the tissue phantom without causing matrix damage, the looped fiber designs were preferable for stabilized interfaces and reliable sensor performance. When integrated into engineered skeletal muscle tissue and agar phantoms with a similar solidity (**Fig. 4B** and **S20**), the sensor provided comparable responses (ΔR/R_0_ = 0.055 ± 0.009 and 0.047 ± 0.01, respectively, at a ≈7% strain stress). Compressing and tensioning of the phantom block caused a sensor response with negative and positive relative resistance values, respectively (**Fig. 4D-E**, **S21**), and cyclical phantom tensioning resulted in a corresponding cyclical sensor signal (**Fig. 4F**), suggesting that the sensor could reliably operate within mechanically compliant matrices.

**Figure 4.**
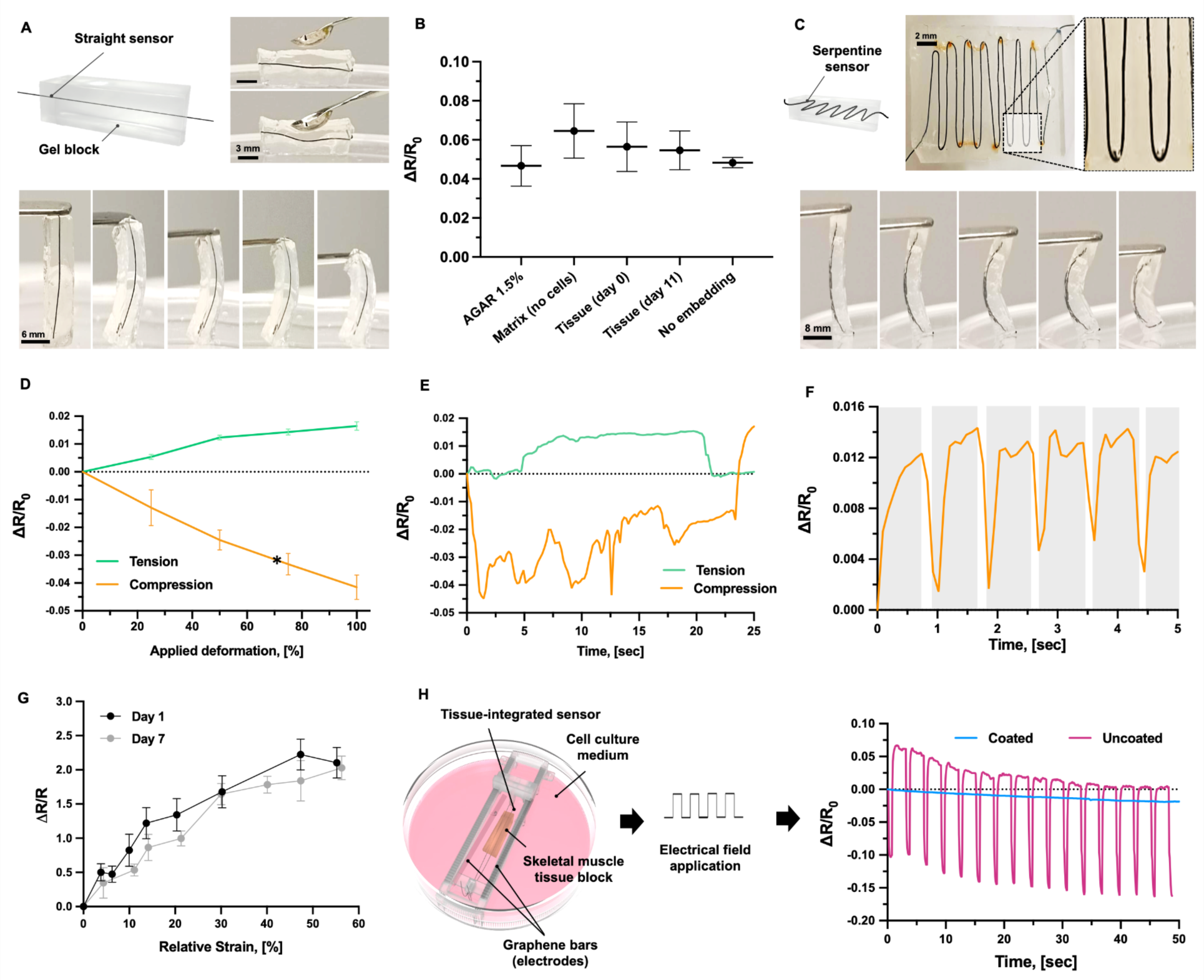
Sensor’s mechano-compliance and operation in an electrical stimulation environment. (**A**) Scheme and optical pictures of the gel-integrated fiber aligned along the longitudinal axis of the gel block, which follows the gel block deformation under mechanical compression. (**B**) relative resistance variation of a strained fiber embedded in various soft matrices. (**C**) Scheme and optical pictures of the fiber integrated into the gel block as arranged in a serpentine design and deforming undergoing mechanical compression. Relative resistance variation of the serpentine-designed tissue integrated fiber over percentage applied deformation (**D**) and over time (**E**). (**F**) Relative resistance variation upon application of cyclic tensile stimuli with 1 Hz frequency. (**G**) Strain-dependent variation of the relative resistance of the tissue-integrated fiber measured at days 1 and 7 after biofabrication (black and gray, respectively). (**H**) Variation of the relative resistance (ΔR/R_0_) of coated and uncoated sensor fibers in the presence of an applied electrical field within a cell culture environment, showing signal interference on the uncoated fiber.

We then assessed the sensing performance of the fiber after long-term exposure to conditions required for cell culture that simulate *in vivo* environments (**Fig S22-23**). After incubating the sensor in cell culture media at 37℃ for one week, the relative resistance response of the uncoated fibers switched from positive values with a strain-dependent increasing profile (**Fig. 2F**) to a non-monotonic response (**Fig. S22**), suggesting that, in a biological environment, possible deterioration of the fiber occurred without the coating. In contrast, the coated fiber retained a response with a monotonic trend (**Fig. S22,** left), which could be useful to sense micromotions from bio-actuators (0-2% stress range). We also exposed the fibers to an active cell culture environment, in which cell bioactivity affected the medium’s composition (**Fig. S24**). We found that while the strain-dependent variation of the relative resistance of the coated fiber matched the strain response measured in air (**Fig. 2F**), the uncoated fiber showed an unreliable performance in tissue. Thus, the coating contributed to preserving the fiber and its function. When embedding the sensor within an engineered skeletal muscle tissue construct, we observed that the strain-dependent relative resistance variation measured at two points in time (days 1 and 7) did not significantly differ (**Fig. 4G**), thus indicating that the fiber could retain its functionality even when embedded in a soft cell-populated matrix that evolved into engineered tissue. Thus, the sensor functionality was compatible with tissue formation and cell culture conditions.

Finally, to test the effectiveness of the coating in insulating the sensing fiber, we exposed the tissue-embedded sensors to a 3Hz electrical pulsated field within a cell culture medium bath (**Fig. 4H**). As shown by the sharp fluctuations of the uncoated sensor response when the electrical pulses were applied, the electrical field interfered with the sensor response of the uncoated fibers. In contrast, the coating successfully shielded the fiber from the applied voltage.

### 2.5 Sensing mechanical deformations of actuated skeletal muscle tissue

We assessed if our biocompatible, soft sensor could help detect the mechanical stress produced by the active contraction of bio-actuators. Therefore, we combined the coated fibers with a ring-shaped, mm-scale skeletal muscle tissue construct that contracted in response to electrical stimulation. The fiber was looped in the muscular ring to reach a co-axial positioning with the bio-actuator, which exploits the tensioning responsivity of the sensor (**Fig. 5A**). We constrained the tissue construct on one side. The fiber was straightened before starting the sensing experiments. We allowed the muscle tissue construct to reach full maturation of myofibers to maximize its output force (**Fig. 5B**).

**Figure 5.**
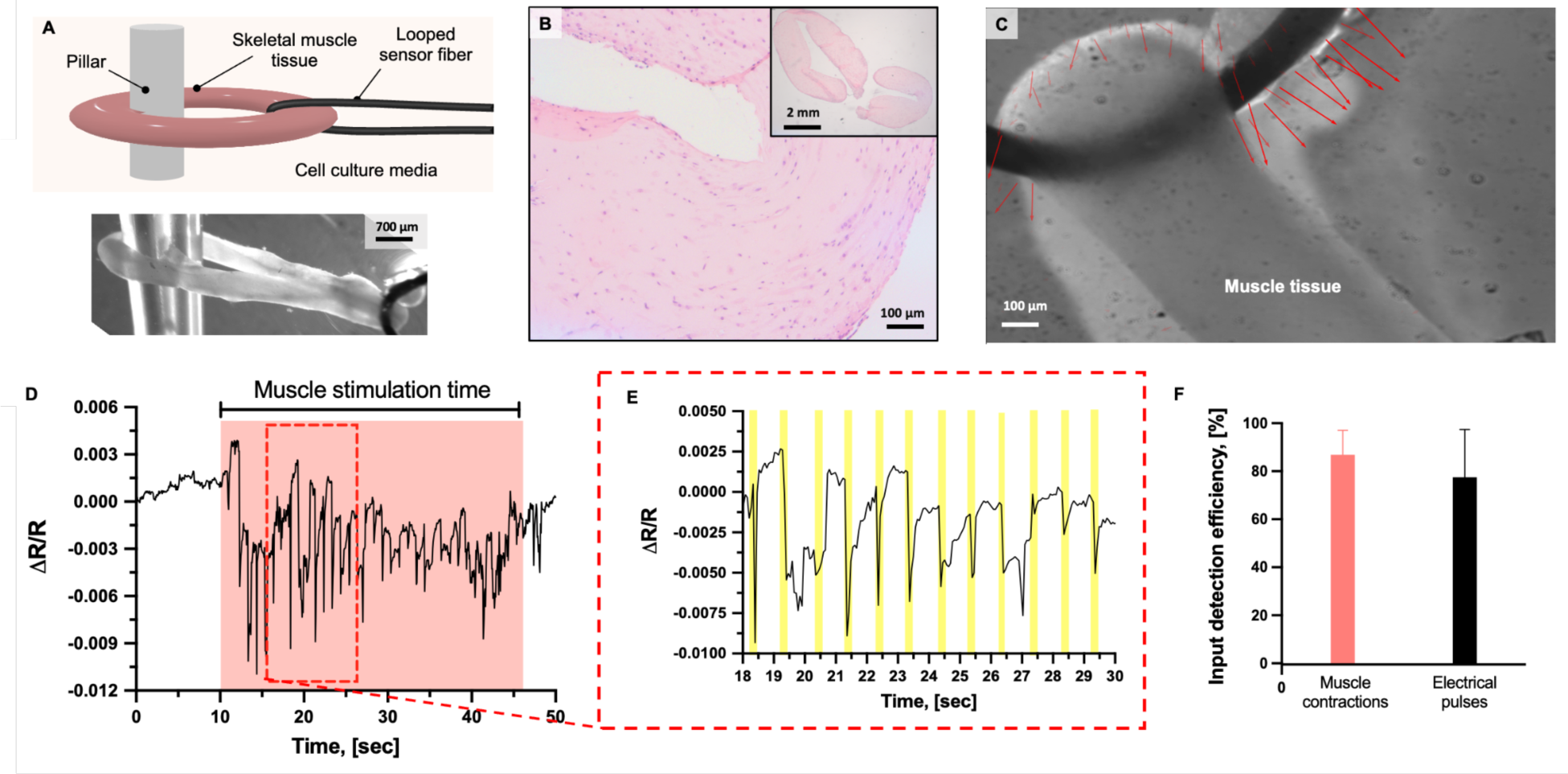
Sensor response of the induced contraction of bio-actuators. **(A)** Scheme of the experimental setup design with a sensing fiber looped in a mm-scale, ring-shaped skeletal muscle tissue construct (top) and optical microscopy picture of it (bottom). (**B**) H&E staining of the skeletal muscle tissue, revealing active, functional myofibers. (**C**) Frame from optical flow analysis of motion video, showing contraction motion of the bio-actuator in the direction of the arrows. (**D**) Relative resistance variation over time of the fiber looped in the bio-actuator during its electrical stimulation. (**E**) Sensor response patterns show peaks of relative resistance variations occurring and matching the stimulation frequency. (**F**) Percentage detection efficiency was calculated by associating relative resistance peaks with microscopically observed muscle contractions and delivery of electrical pulses.

Upon electrical stimulation, as visualized by optical flow analysis (**Fig. 5C**), the construct produced a twitching motion that matched the electrical stimulation pulse scheme provided in the frequency domain of 0.5-3 Hz. When stimulating the engineered muscle tissue with 1 Hz direct current pulses (1% duty cycle), the relative resistance variation was recorded as the fiber moved from the equilibrium state to the active actuation (**Fig. 5D**). The time point when the bio-actuator started its contraction could be identified in the recorded signal, as a variation of relative resistance (≈0.024±0.007) greater than baseline noise was detected between the “actuation phase” signal with the baseline signal before the pulse was applied. Signal perturbations occurred in a confined value range (0-0.01), mostly in the negative value space (**Fig. 5D** and **E**).

We attributed the small range to the very low tensile forces exerted on the fiber by the muscle’s contraction, as shown by the tensile characterization at low strains (**Fig. 2G-H**). The signal of the sensor response peaked with a similar frequency to the applied electrical pulsation pattern (1 Hz, **Fig. 5F**). By analyzing the video recording of the bio-actuation performance, we calculated that the fiber had a high input detection efficiency, as almost 90% of the observed contractions could be associated to measured peaks. However, the number of sensor peaks was lower than the number of electrical pulses applied (≈75%, **Fig. 5F**), potentially due to the lack of muscle responsiveness to a few stimuli and, thus, a minor amount of mechanical inputs acting on the fiber.

### 2.6 Closed-loop control of the bio-actuator

To understand whether the sensor could be used to control the behavior of a bio-actuator, we built a platform in which the sensor was integrated into the maturing skeletal muscle tissue. After a 24-hour growth period, a ring-shaped bio-actuator was looped at its two extremities with two sensing fibers that were wrapped around pillars (**Fig. 6A-B** and **S25-27**). The fibers equally tensioned the tissue to deliver a balanced mechanical stress on the forming skeletal muscle tissue that promoted cell alignment (**Fig. 6A-B**). During tissue development, soft tissue grew around the sensor, thus forming a continuous tissue-sensor interface (**Fig. 6A**). In this configuration, the sensor-active fiber portion corresponded to two four-mm-long segments for each bio-actuator’s extremity (**Fig. 6C**). After 11 days of myogenic differentiation, the bio-actuator responded to electrical stimulation with rhythmic contractions that matched the delivered pulse frequency (1Hz, **Fig. 6D**).

**Figure 6.**
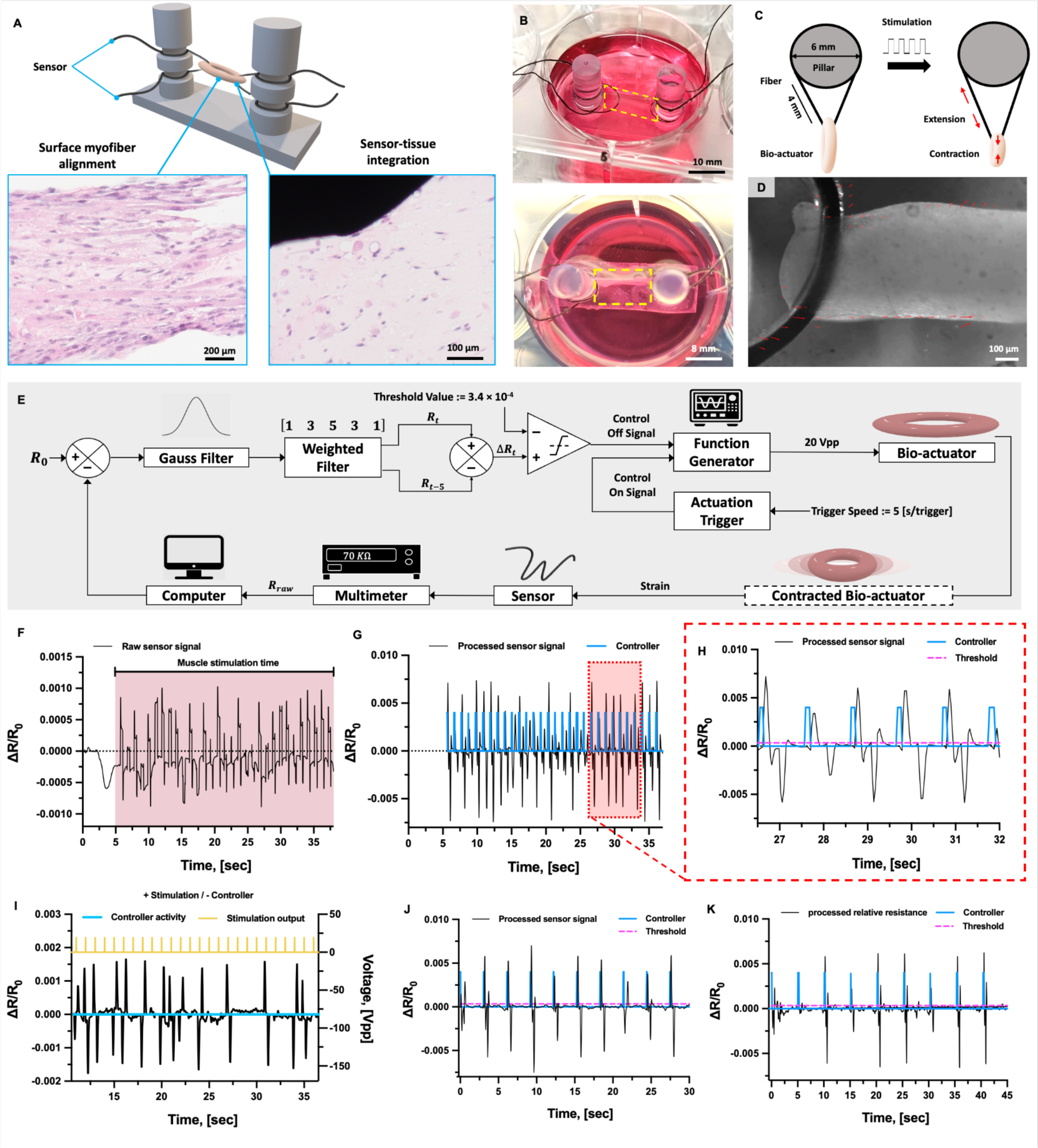
Closed-loop bio-actuator co-developed with the sensor fiber. **(A)** Scheme of the experimental setup design in which the bio-actuator was looped in two sensor fibers, and H&E histological staining of the muscle tissue (left) and tissue-fiber interface (right). (**B**) Side (top) and top (bottom) view of the platform with the tensioned bio-actuator (yellow dashed box). (**C**) Top-view schematics of the setup elements. (**D**) Frame from optical flow analysis of motion video, showing contraction motion of the bio-actuator in the direction of the arrows. (**E**) Schematics of the closed-loop control on the contracting bio-actuator that creates a mechanically-informed actuator’s on/off switch. (**F**) Sensor response of the bio-actuator under 1Hz control trigger. (**G**) Processed sensor signal (black line) and real-time behavior controller (blue line) in a closed-loop system. (**H**) Detail of the sensor signal and controller, showing reliable controller operation. The threshold value is shown (dashed pink line). (**I**) Processed sensor signal (black line), controller’s activity (blue line), and stimulation output (yellow line) in an open-loop system, where no controller is applied. Sensor response of the bio-actuator and controller behavior under variations of the control paradigm with triggers set at 3 (**J**) and 5 (**K**) secs.

For closed-loop control, we designed a controller that triggers a stop to electric field application when measuring a resistivity change above a predetermined threshold (**Fig. 6E, S25-27**). By analyzing the bio-actuator’s contraction as revealed by the sensor as peaked variations from the baseline sensor signal (**Fig. 6F**), we set the threshold value to correspond to a strong contraction of the bio-actuator (**Fig. 6G**). To study the closed-loop system functionality, we gave the controller a periodical trigger to turn on the electric field application at different intervals. After sensing the contraction of the bio-actuator, the controller would then turn off the electric field until the next trigger (**Fig. S28**). By comparing the processed sensor response and the actual controller behavior (black and blue lines, in **Fig. 6G-H**, respectively), we observed that the system responded reliably with 100% of expected controller responses (**Movie S1**). The stimulation output in the open-loop system was not affected by the bio-actuator’s response (**Fig. 6I**), and the control system could not affect the stimulation output in the absence of applied stimuli (**Fig. S29** and **Movie S2**). Moreover, the system successfully performed with different triggers (1, 3, and 5 sec) and different thresholding (**Fig. 6J-K**, and **Fig. S30**).

The closed-loop system could be used to program bio-actuators’ stimulation programs that are tailored to the tissue responsivity (**Fig. 7A**). At an early stage of the muscle tissue maturation (day 7), the bio-actuator’s response to electrical stimulation was observed (**Fig. 7B**, **Movie S3**). The signal intensity of the sensor response differed from the one measured at the end of the maturation time (day 11, **Fig. 7C** and **D**), suggesting this platform could be used to study the tissue development process and the mechanical forces involved in it. The closed-loop system could regulate the electrical stimulation used as a training tool during tissue development (**Fig. 7E**), or control the bio-actuator’s performance by adapting to the specific actuator’s response to maximize force output and synchronization while minimizing the unnecessary exposure of the tissue to electrical fields (**Fig. 7F** and **G**).

**Figure 7.**
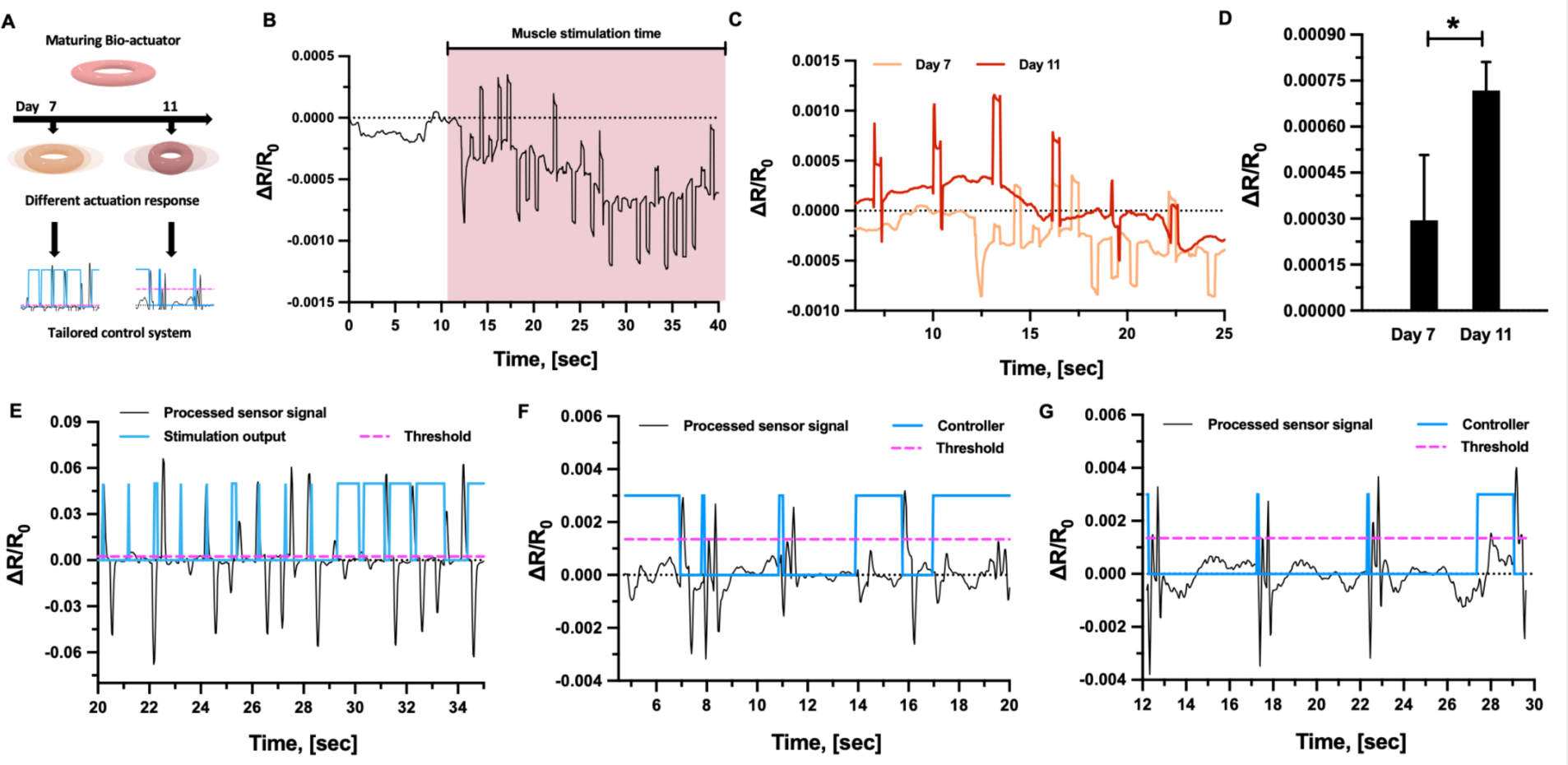
Performance exploration with a closed-loop bio-actuator. (**A**) Schematics of tailored stimulation as programmed on the different bio-actuator’s responses. (**B**) Raw sensor signal from a bio-actuator’s response at day 7. (**C**) Time-dependent sensor signal from the same bio-actuator tested on days 7 and 11. (**D**) Sensor signal intensity from the bio-actuator at the two-time points as calculated from the spike height from the raw signal (analyzed time interval: 7.5-25 secs). (**E**) Closed-loop system applied to sensor signal acquired on day 7. Control system adaptation to different bio-actuator’s responses: processed sensor signal (black line) and real-time behavior controller (blue line) in closed-loop actuation (thresholds: 0.0013; trigger: 3 secs) in response to less (**F**) and more evident (**G**) bio-actuators’ response patterns.

## 3. Discussion

Bio-hybrid robotics has emerged as a frontier area of robotics looking at soft, renewable, and intelligent materials for compliant motion realization (*1*). Various bio-hybrid systems relying on different mechanisms of activating muscle tissue have been demonstrated, including electric field application (*9*), wireless powering (*22*), and optogenetics (*23*). However, all of the previously demonstrated examples rely on external motion control triggered by an observer. Achieving autonomous motion sensing and control is a fundamental challenge to unlock the potential of future bio-hybrid robots. With our work, we took the first step towards intelligent autonomous motion by demonstrating closed-loop control of bio-actuator contraction, achieved by integrating a soft and biocompatible fiber-shaped sensor into the bio-actuator. Our bio-actuator can sense and subsequently act on its contraction state.

To realize our proprioceptive bio-actuator, we developed a novel tissue-integrated mechanical sensor. Mechanical sensors are garnering relevance in biomedical engineering due to their ability to detect and measure mechanical forces and deformations in biological systems, biomedical models, and biometrical devices (*24–26*). A mechanical sensor embedded into fragile bio-systems and combined with simple readout methods would allow us to monitor the structural stresses, control the motion, and understand tissue repair and formation processes (*27*, *28*). However, for use in bioengineering applications, the component materials, designs, and manufacturing approaches of mechanical sensors have to be optimized to improve the performance of the sensors (*i.e*., sensitivity, durability) within biological environments that feature complex architectures, soft materials, liquid milieu, and activity of living systems (*i.e*., cells) (*29*, *30*). Developing biocompatible, reliable, and minimally invasive sensors poses a significant challenge, limiting their application to natural or engineered biological systems (*31*, *32*).

Recent advances in micro- and nanotechnology have enabled sensors with flexible, miniaturized, and less invasive designs, thus creating new opportunities for direct contact and implantable sensor technologies (*33*). Various mechanical sensor types have been used for biomedical engineering applications including piezoelectric, capacitive, and piezoresistive sensors. Due to their versatile strain-sensing abilities, easy readout, and low energy requirements, piezoresistive materials have been widely investigated for robotics, biomedicine, and other applications (*e.g.*, for biometrics and prosthetics), and enable wearable electronic devices, electronic skin, and flexible sensors, and implantable interfaces (*34*, *35*). Despite their potential for sensing technologies, using piezoresistive composites for biomedical sensors remains underexplored. To perform in biomedical devices and integrate into living tissue, piezoresistive sensors have to feature mechanical compliance with tissues, sensitivity (i.e., high Gauge Factor), and monotonic response, particularly under dynamic conditions. Further, the sensor must be electrically insulated from the conductive tissue environment and sensor readouts should have low piezoresistive hysteresis and viscous drift and be stable over weeks in culture.

To engineer a sensor that could meet all of these criteria, we selected piezoresistive composites based on carbon black (CB)-enriched styrene-ethylene-butylene-styrene (SEBS) thermoplastic elastomers, which display high electrical conductivity, good mechanical properties, and safe manufacturing strategies (*36*). Importantly, both CB and SEBS have been individually used for tissue engineering and other biomedical applications, demonstrating they can be safely incorporated with living materials (*37–40*). However, the combination of CB and SEBS to form mechanosensors for optimized bio-integrated functionality has never been investigated. Our work demonstrated that these materials can combine with living tissue that is engineered *in vitro*, operate at high sensitivity, and retain long-term functionality. We maximized the softness of our material and shaped it into a fiber design of a small diameter (<0.25 mm) to enable versatile tissue-integrated designs and stabilize the bioelectronic interface between tissue and sensor. An insulation layer around the fiber was necessary to preserve the response signal from disturbance of other electrical inputs present in a cell stimulation setup, but also served to protect the structural integrity of the sensor over time. Our engineered tissue matrices and piezoresistive sensors can be assembled and grown together, without any negative effect on engineered tissue formation or decay of sensor functionality.

We also aimed to demonstrate intelligent control of bio-actuators’ motion. Current bio-hybrid systems lack system integration, which can be achieved by associating actuation with sensing and control strategies to generate operational autonomy. Currently, understanding the bio-actuators’ position and quantifying their movement are mostly performed by external observers via optical methods and dynamical analysis of recorded videos. For example, Iuliano et al. developed a tissue engineering platform that integrates optical fiber-based sensors to measure via light interferometry the displacement of a flexible cantilever induced by ring-shaped skeletal muscle tissue (*41*). Optical approaches have limited applicability to robotic systems due to illumination and imaging requirements. In contrast, our piezoresistive sensor can operate in any light condition and without imposing design constraints on the system. In addition, the tissue intimate contact allows one to detect tissue contractions expressed through minimal forces with high accuracy and spatiotemporal resolution.

A few strategies that enable integrated readouts of bio-actuator contraction without external observation have been explored. For example, Kim et al. proposed to measure the bio-actuators’ contraction forces with a mechanosensor made of liquid Galinstan embedded within soft elastomeric substrates (*42*). Though the authors described the simulated performance of the sensor, the actual integration of the sensor into the tissue was not attempted, and no feedback control on the tissue functionality setup was shown. The lack of demonstrated tissue functionality could be attributed to the presence of rigid components in the sensor which hamper tissue integration and cause strain shielding effects. In contrast, our flexible sensor fiber is biocompatible and consistent in its design, can easily integrate with both the engineered tissue and the reading interface, and can operate on a cultured living muscle.

Zhao et al. have also recently reported a flexible mechanosensing device using a microscale strain sensor that can be used to measure mm-scale engineered muscle contractions (*43*). The developed device is biocompatible, flexible, and was demonstrated to read optogenetic muscle contractions over time. However, the geometry of the device is limited to a tissue-on-a-chip design as the strain sensors are cleverly integrated into a planar system designed as a holder for a ring-shaped muscle, and force contraction sensing comes from the deformation of the holder. Therefore, this device has a strong limitation on the achievable geometric complexity of the bio-actuators and does not integrate with tissue. Further, the device functionality was proven only with optogenetic control of muscle contraction, and not with electrical stimulation of wild-type muscle cells. Our insulated fiber-shaped sensor is functional in baths for cell electrical stimulation and offers the freedom to design tissue interfaces in 3D geometries inside soft matrices. Finally, our sensor is completely based on soft materials, without the integration of any rigid or metallic components. These features open perspectives to realize complex sensor arrangements for implantable devices, 3D tissue models, and future motile bio-robots with portable designs.

Finally, we used our sensor to design a first-in-class proprioceptive bio-actuator. We generated an artifacts-free mapping between the muscle contraction and relative resistivity change in the fiber that confirmed the detection of bio-actuators’ deformations, which typically occur as micromotions. This mapping enabled the design of a closed-loop controller, which then realized the first, auto-determined behavior in a bio-hybrid actuator. In our closed-loop demonstration, we showed that a tissue contraction event could trigger a switch-off action of a controller with a high correspondence between the sensor signal peaks and the controller output turn-off points. In our sensor-integrated developing bio-actuators, the control system enabled autoregulated electrical stimulation that could serve for tissue maturation under training or operation in bio-hybrid robots. We acknowledge that the contractile response of our sensorized, tensioned bio-actuators matched the performance expected from state-of-the-art bio-actuators.(*1*) As the ability to integrate sensors throughout the differentiation process contributes to controlling the tissue maturation process, we expect our system to participate in improving the bio-actuators’ contractile ability, as the next challenge to be addressed in bio-hybrid robotics. Intriguingly, as our sensor could combine with other chemo-electroactive cells (such as neurons), wider applicative horizons in bio-mechanics and bioelectronics (such as multi-functional organ-on-a-chip devices) are possible, which would benefit from the sensor’s elastomer-based composite nature that combines a soft mechanical profile with the potential to cover a wide sensing range at low fabrication costs (*34*).

Overall, our data represent a stepping stone for the generation of fully integrated biohybrid robots and a new starting point to research portable designs and onboarded multifunctionality in organic machines. We foresee that the next efforts will look at defining strategies for energy-efficient control systems, as well as the realization of more complex reaction schemes. For example, one next challenge will be the creation of adaptive control signals that increase the lifespan of bio-actuators’ functionality by adjusting the stimulation paradigms to consider the natural deterioration or functional decay of the engineered tissue. Despite many advances in the field of bio-hybrid robotics, bio-hybrid systems are largely underdeveloped technologies and their performance cannot compare with traditional robotics under various metrics (*1*).

Our concept to sensorize bio-actuators will enable biohybrid robotics to overcome the challenge of full operative integration, inspire novel forms of biological machines capable of intelligent motion, and thus unlock the potential of biological robots, which, as endowed with autonomous operativity for real-life applications and will further express the unicity of their living component, through adaptability, self-healing, biodegradability, efficiency, and mechanical compliance. In a wider sense, the reported results can inspire technologies with direct application to pathophysiological 3D tissue models, micro-physiological systems, pharmaceutical research, and bio-hybrid system engineering.

## 4. Materials and Methods

### 4.1 Thermoplastic processing of the sensor fibers

Implant grade styrene-based tri-block co-polymer (TPS) was donated by Hexpol (Malmö, Sweden) in Shore hardness 20A, 40A and 50A. Carbon Black Ensaco 260G was obtained by Imerys (Paris, France). TPS and Carbon Black were mixed in 1:1 mass ratio, using the torque rheometer Polylab OS from Thermofisher (Karlsruhe, Germany). The temperature during mixing was 180°C. After the melt-mixing step, the composites were extruded (180°C) in the form of fibers with a diameter of 0.2 mm, using the capillary rheometer RH7 from Netzsch (Selb, Germany). The fibers were pre-strained and let relax, and their final diameter was confirmed with the optical microscope Zeiss Stereo Discovery (Carl Zeiss Microscopy, Jena, Germany).

### 4.2 Dip coating of the sensor fibers

For the dip coating of the fibers, a liquid silicone rubber (LSR) EcoFlex 00-30 from Smooth-On (Macungie, PE, USA) was used in a 1:1 ratio. The fibers were coated by dipping them twice into the silicone rubber and leaving them to dry on aluminum foil.

### 4.3 Mechano-electrical characterization of the sensor fibers

The mechano-electrical testing was performed with the tensile testing machine Zwick Roell Z005 (ZwickRoell, Ulm, Germany). A 200 N load cell was used. The fibers were clamped with a pressure of 4 bar applied by pneumatic clamps. The electrical resistance was recorded during the tensile testing with a source meter (#2450, Keithley, Cleveland, USA). A voltage of 1 V and a sampling frequency of 10 Hz were used to characterize the fibers by tensile testing and 1000 Hz for the fibers integrated in the biorobots. A gauge length of 50 mm and a strain rate of 200 mm/min were used for the tensile test investigation. The relative resistance R_rel_ was calculated by:

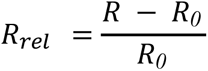

where R is the measured resistance and R_0_ is the measured resistance at the beginning of the tensile test. Data were presented as mean values ± standard deviation with sample size (*n* = 5).

### 4.4 Scanning electron microscopy

The samples were fixed with 2.5% (v/v) glutaraldehyde in PBS solution at room temperature for 30 min and then rinsed three times with PBS. Then, the samples were dehydrated in an ascending series of ethanol solutions and dried in a critical point dryer (Tousimis 931). For conductivity, the samples were sputter coated with 5 nm Pt/Pd (Safematic CCU-010). The examination was done in a JSM-7100F JEOL SEM at 3 kV by secondary electron detection.

### 4.5 Functionality of the fiber embedded in soft matrices

To generate the tissue-mimicking phantoms, we cast various polymeric matrices into rectangular-shaped molds (size: 30×20×10 mm). Agar matrices were prepared by gelating different water solutions of agar (#A1296, Sigma Aldrich) at various w/v concentrations (from 0.5 to 2%). A matrix with no cells was prepared by mixing Matrigel basement membrane matrix (Corning, New York, USA) and Collagen (Cosmo Bio, USA) at 1:1 ratio. The same matrix was tested immediately after myoblasts addition and after several days of tissue development. An analog Shore Hardness tester (HB0 100-0, Sauter, Balingen, Germany) was used to measure the shore hardness at room temperature 2 hours after preparation. Two embedding pre-tensioned designs were studied, with the fiber placed in straight or serpentine poses. The surface adhesion and the flexibility of the fiber within stressed phantoms (1.5% w/v agar, size: 25×10×8 mm) under manual mechanical compression were shown with representative optical pictures.

### 4.6 Cell culture

The murine myoblast cell line C2C12 was obtained from the American Type Culture Collection (ATCC, Manassas, VA, USA). Cells were cultured in monolayer at 37 °C in a 5% CO_2_-containing humidified atmosphere in a complete growth medium (GM), consisting of Dulbecco’s modified Eagle’s medium (DMEM, #D6429, Sigma–Aldrich) supplemented with 10% (v/v) heat-inactivated fetal bovine serum (FBS, #F7524, Sigma–Aldrich), 2 mM glutamine, 100 U mL^−1^ penicillin, and 100 μg mL^−1^ streptomycin (all from Thermo Fisher Scientific, Switzerland). To induce cell myogenic differentiation, a differentiation cell culture medium (DM) was used, which consisted of DMEM containing 10% horse serum (Gibco), 1% penicillin-streptomycin, IGF-1 (50 ng/ml; Sigma-Aldrich), and ACA (1 mg/ml; Sigma-Aldrich). To prove the biocompatibility of the sensor components, other cell types were used.

### 4.7 Engineered skeletal muscle tissue fabrication and fiber integration

Block-shaped skeletal muscle tissue constructs were prepared by mixing Matrigel basement membrane matrix (Corning, New York, USA) and Collagen (Cosmo Bio, USA) (ratio 1:1) with 1×10^8^ cells/mL and manually cast in a rectangular block-shaped mold. The constructs were left in a cell incubator for 30 min, unmolded, and cultured in GM and DM for 3 and 6 days, respectively. Cells suspended in a GM were mixed with Matrigel and fibrin (Sigma-Aldrich) to fabricate ring-shaped constructs. The mixture was cast in a planar ring-shaped mold and cultured for growth and differentiation medium for 3 and 10 days, respectively, as described elsewhere (*44*).

### 4.8 Live/dead staining in 3D constructs

We assessed cell viability in the constructs using Live/Dead™ staining following the manufacturer’s instructions (Thermo Fisher Scientific; #R37601). Calcein and propidium iodide (λ_ex_=488 and 561 nm, respectively) were imaged on a confocal microscope (Zeiss LSM 780 Airyscan, Zeiss AxioObserver.Z1, ScopeM). Imaging analysis was performed on at least three images from each analyzed sample feature (≥ 3 samples/condition; ≥ 3 experimental replicates).

### 4.9 Cell viability tests and cell growth in 3D constructs

Resazurin assay (AlamarBlue, #R7017, Sigma-Aldrich) was used to evaluate the biocompatibility of the fiber materials and cell growth within the scaffolds, following the manufacturer’s instructions. All cells were seeded in 96 well plates at an initial cell density of 5×10^3^ cells/cm^2^, and then incubated with different amounts of fiber materials (from 0.1 to 100 mg/ml) for variable durations (from 6 hours to 14 days). For 3D constructs, AlamarBlue reagent was added to the media. The fluorescence signal (λ_Ex/Em_ = 530/590) was measured with a Synergy H1 microplate reader (Biotek). Fluorescence intensity values were corrected for the background control (culture medium with resazurin).

### 4.10 Immuno-fluorescence in 3D constructs

After *in vitro* culture, the constructs were fixed overnight in a 4% Paraformaldehyde solution at 4℃, rinsed in PBS (5 min, 3×), and stained for f-actin (AlexaFluor 488 phalloidin, #R37110, Invitrogen), MyoD (primary Anti-MyoD Antibody, #ZRB1452, Sigma-Aldrich, and an anti-rabbit secondary antibody coupled with Rhodamine #SAB3700846, Sigma-Aldrich), DAPI (4′-6-diamidino-2-phenylindol) and Myosin Heavy Chain (Myosin 4, eFluor™ 660, Clone: MF20, Affymetrix eBioscience™). All images were taken using a confocal microscope (Zeiss LSM 780 Airyscan, Zeiss AxioObserver.Z1, ScopeM) and analyzed in Fiji ImageJ. The number of nuclei within MyHC-positive myotubes was counted in 5 fields per sample, and used to calculate the fusion index as a ratio against the total number of nuclei. The images were randomly selected from at least 3 regions from each sample, and the experiments were replicated thrice.

### 4.11 Immunohistochemistry on tissue sections

After *in vitro* culture, the constructs were fixed overnight in 4% Paraformaldehyde solution at 4℃ before they were embedded in paraffin, and sections of 4.5 μm thickness were cut with a microtome (Microm, HM430, Thermo Scientific). The sections were stained with Hematoxylin-Eosin (#GHS116 and #HT110116, respectively from Sigma Aldrich) and imaged with a light microscope (Olympus CKX41, Olympus Schweiz AG).

### 4.12 Sensing bio-actuator’s contractions and motion analysis

Six-well plates were coated with 2% agar (#A1296, Sigma Aldrich) to form a 0.5 cm-thick layer, and the muscle tissue ring-shaped constructs were transferred into them with DMEM. The coated fiber was looped inside the muscle tissue rings. A pillar was inserted through the hole of the muscle ring into the agar bed in a vertical position. The fiber was anchored to the opposite side of the culture well and tensioned. The fiber extremities were attached to a source meter (#2450, Keithley, Cleveland, USA), and electrodes were connected to a function generator (#DG812, Rigol, Beijing, China). A custom-made holder with two graphene electrodes was inserted into the culture medium, and electrical cell stimulation was performed by applying square pulses with 1Hz frequency at 4.5 V/cm (10 V; electrode distance: 2.2 cm) and 1% duty cycle. The experiment was recorded using a Zeiss Stemi 508 stereomicroscope and a Basler ace2 camera at 25 fps.

### 4.13 Optical flow analysis of tissue contraction

We used an open-source (OpenCV) optical flow algorithm with Gunnar Farneback’s algorithm to visualize tissue contraction (*45*, *46*). Videos are analyzed frame-by-frame to provide a dense pixel-wise velocity between subsequent frames. For visualization purposes, arrow lengths were scaled by a factor of 100 and shown as a uniform sampling of 1% of the pixels.

### 4.14 Control system

To verify if the sensor system could be used in a closed loop controller, we analyzed the relative resistivity change measurements on the fiber obtained by applying square pulses with 1Hz frequency at 4.5 V/cm (10 V; electrode distance: 2.2 cm) and 1% duty cycle. Three Gaussian *N*(μ, 1) filters were applied sequentially to the original signal to decrease noise. The filtered signal was then processed with a simplified thresholded Canny edge detector function to determine the presence of significant minima corresponding to the maximum contraction stages of the bio-hybrid setup **(Fig. S26)**. Having verified the capabilities of the measured signal, a threshold closed-loop controller was realized **(Fig. S27)** by feeding the data on an update loop as it became available and performing real-time processing on past measurements. The controller was configured to trigger a simulated signal at the desired frequency (commonly 1 Hz) and analyze the rate of change of relative resistivity in the fiber to determine the shut-off point on each trigger cycle. The turn-off evaluation was performed on every measurement update (sampled at 16 Hz) and realized by applying a Gaussian *N*(μ, 1) filter on all the raw available measured data, followed by a weighted median filter (w = [1, 3, 5, 3, 1]), and an estimation of the rate of change relative to the fifth prior measurement.

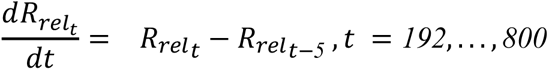

The rate of change is then evaluated with a predefined threshold set to 0.0013 to differentiate between true contractions and noise. The threshold was adjusted where necessary as dictated by the maturation state of the muscle tissue, as well as the strength of the applied voltage.

### 4.15 Data analysis and statistics

Data were analyzed with the Graph Pad Prism 9 Software. All variables are expressed as mean ± standard deviation (SD). Data were acquired from at least three independent experiments and three technical replicates unless otherwise stated. To assess statistically relevant differences between the two experimental groups, the *t*-test was used (P < 0.05 and P < 0.001 are expressed as * and **, respectively). A general linear two-way ANOVA test was used to verify the hypothesis whether there were changes in various parameters over time among the experimental groups and to identify relevant variations among several experimental groups.

## Acknowledgments

We thank Mike Michelis for his help in implementing the optical flow analysis. We thank the members of the DBM Histology Core Facility (University of Basel) for their support and/or the execution of the experiments. The authors also thank Anne Greet Bittermann from the Scientific Center for Optical and Electron Microscopy (ScopeM) of ETH Zurich for acquiring images at the SEM. We would like to thank Louisa Marie Eckey from Empa Dübendorf for the thermoplastic extrusion of sensor fibers.

## Author contributions

All authors planned the experiment and contributed to the analysis of the data and discussion. M.F., A.B., and A.G. performed and supervised the experiments, and wrote the manuscript.

## Data and materials availability

All data are presented in the paper and/or in the Supplementary Materials. Contact M.F. for additional information.

