## Supplementary Information for "Integrated Closed-loop Control of Bio-actuation for Proprioceptive Bio-hybrid Robots"

**INDEX**

### Preface

In this work, we developed a soft, flexible piezoresistive sensor and integrated it into a skeletal muscle tissue construct serving as a bio-actuator. With this innovation, we demonstrated the first proprioceptive bio-hybrid robot that is informed on its biomechanical state and can regulate its behavior accordingly. Here, we report on supplementary data that complement the data collection shown in the main manuscript. The present document includes additional results describing: the sensor functionality as assessed via mechanoelectrical characterization; the biocompatibility of the sensor on different cell types; and its integration within 2D and 3D cell culture models in the presence of applied electrical fields. Moreover, we show more information about the sensor's ability to reveal micro-motions generated by electrically induced contraction of bio-actuators, and we present our design of a feedback control system that processes the sensors' response data.

### 1. Sensor characterization

To realize our sensor, we studied different material compositions and characterized the sensor's functionality via mechanoelectrical characterization, and we proved its biocompatibility and integration with various 2D and 3D cell culture systems. The fibers with the lowest Shore hardness studied in this work (20A and 40A) displayed a positive slope with a Gauge Factor (GF) of 10 at 10% strain. At 50 % strain, the GF for the 50A fiber and the other two fiber types was 6 and 8.4, respectively. We characterized the strain-dependent stress and the relative resistance variation of the 20A fibers in the presence or absence of coating (**Fig. S1**).

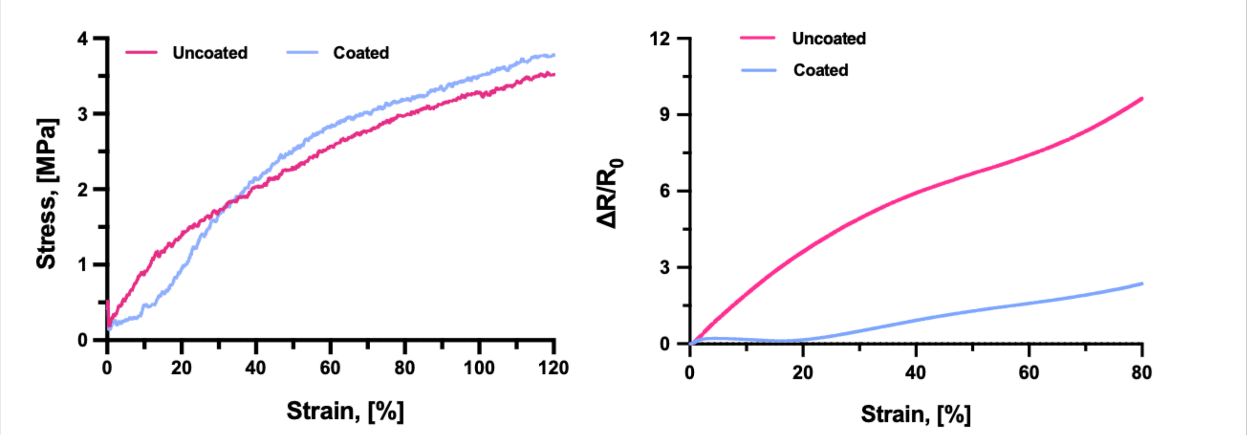

**Figure S1. Strain-dependent stress response of the pre-strained sensor fiber.** Strain-dependent (0-120% strain range) variation of the stress (left) and relative resistance (right) of the coated and uncoated pre-strained fibers (blue and red lines, respectively) in the 0-120% and 0-80 % strain range.

When subjected to dynamic tests with a train of consecutive applied strain stimuli, the fibers demonstrated repeatable stress and resistance responses. We studied dynamic responses to repeated 50% (**Fig. S2**). The coherent behavior between stress and resistance response observed in the dynamical tests proves that the sensor response is reliable.

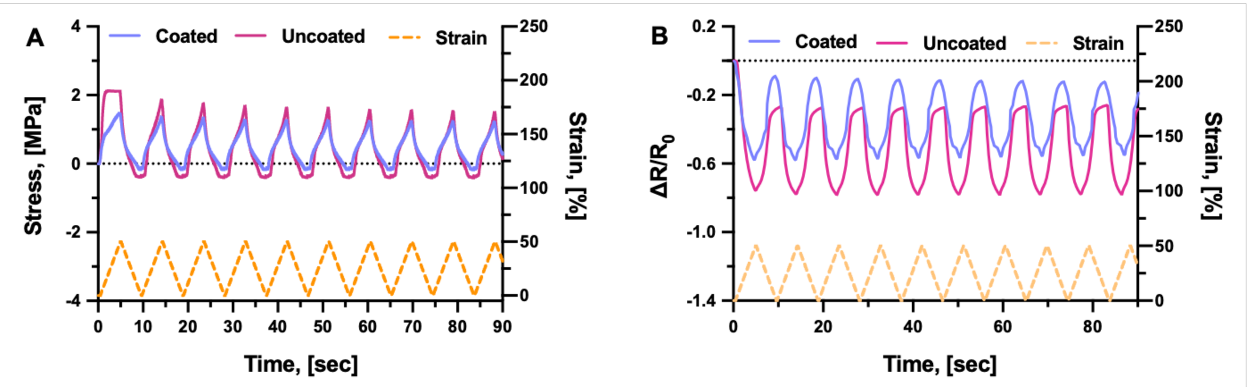

**Figure S2. Sensor dynamic response.** Time-dependent variation of the stress (**A**) and relative resistance (**B**) of coated and uncoated pre-strained fibers (blue and red, respectively) under repeated strain stress applied at 50%.

To understand if our sensor could serve to detect minimal displacements likely caused by small output forces of bio-actuators, we tested the pre-strained fiber in the 0-1% strain ranges, finding that the stress response coherently reacted to the applied strains (**Fig. S3-5**).

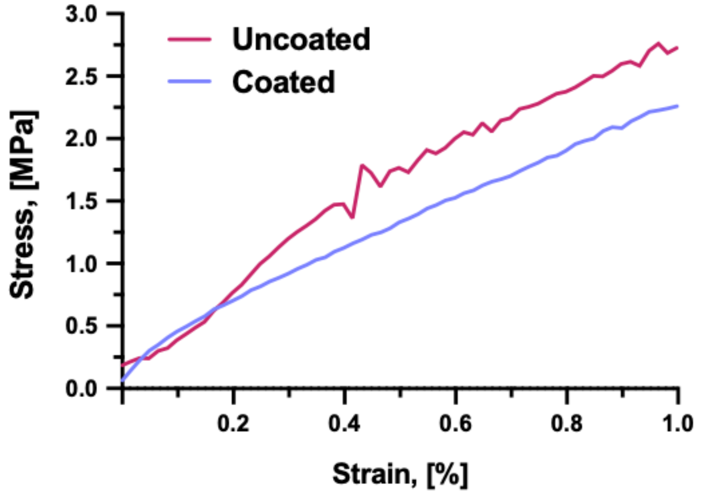

**Figure S3. Strain-dependent variation of the stress response in the low strain range.** Strain-dependent variation of the stress of coated and uncoated pre-strained fibers in the 0-1% strain range.

We observed that the stress-strain slope decreased when coating the fiber with a silicon layer. This observation can be explained by the fact that the silicon layer has a lower Shore hardness, and by the relationship between Shore hardness and Young's modulus explained by Gent in 1958 (1). According to the composite theory (2), the overall Young's modulus of the coated fiber will be a mixture of the fiber and the coating Young's modulus, which takes the volume content of both into account. Therefore, Young's modulus of our coated fibers decreased, as shown in **Fig. S3**.

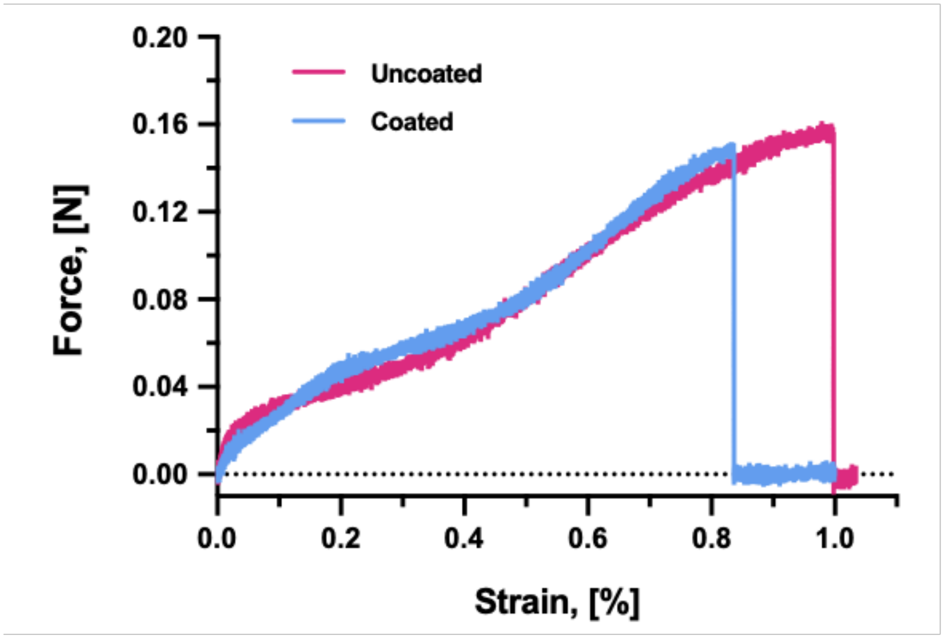

**Figure S4. Strain-dependent applied forces on the pre-strained fiber.** Strain-dependent (0-1% strain range) variation of the tensile forces applied to the coated and uncoated pre-strained fibers (blue and red lines, respectively). The decrease in force is not due to fiber breaking but rather to being pulled out of the holding system.

To understand if the sensor can operate consistently during trains of stimuli occurring during the mechanostimulation of a bio-actuator, we tested the fiber response in dynamic tests with repeated tensile stress for a 0.2% stress (**Fig. S5**).

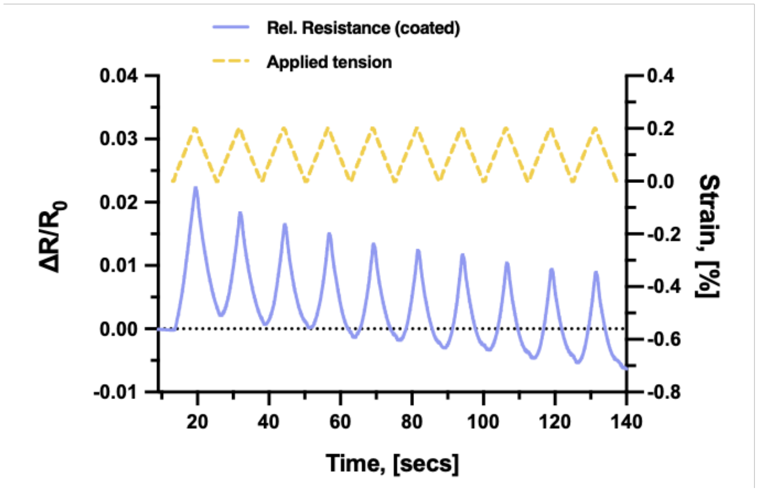

**Figure S5. Sensor dynamic response.** Time-dependent variation of the relative resistance of the coated pre-strained fibers (blue line) under repeated strain stress (yellow dashed line) applied at 0.2%.

We observed that when the fiber was positioned in a slightly loose straight configuration, the baseline signal of the sensor displayed a negative drift (**Fig. S6**). When applying repeated tensile stresses of 0.1 mm total displacement, corresponding to a movement and applied forces in the same value ranges expected for the bio-actuator's activity (*i.e.*, a few mN), peaks in the relative resistance emerged in the negative data space, thus suggesting that sensor signal drift in the negative values would not affect the readout of bio-actuators' motion. Further, the fiber would still work to read signals even from a loose configuration (**Fig. S6** and **S7**).

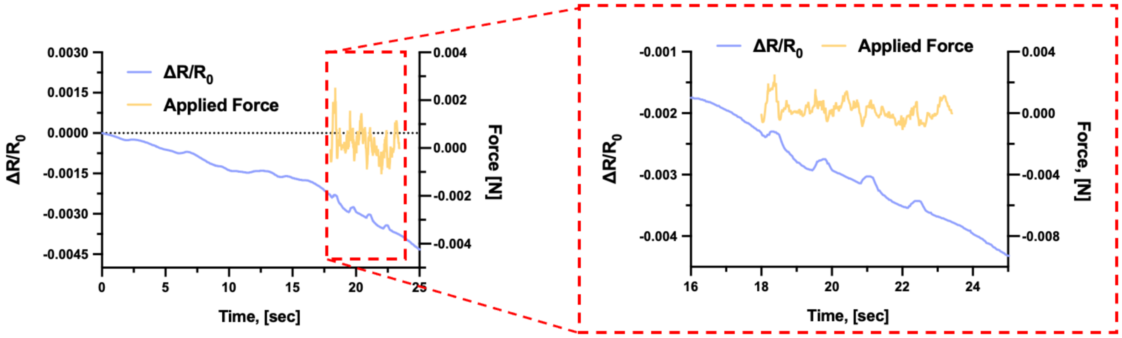

**Figure S6. Sensor baseline from a loose fiber.** Time-dependent sensor response of coated pre-strained fibers showing a negative drift (left). Under tensile stress with a few mN forces, relative resistance in the negative data space is detected (right).

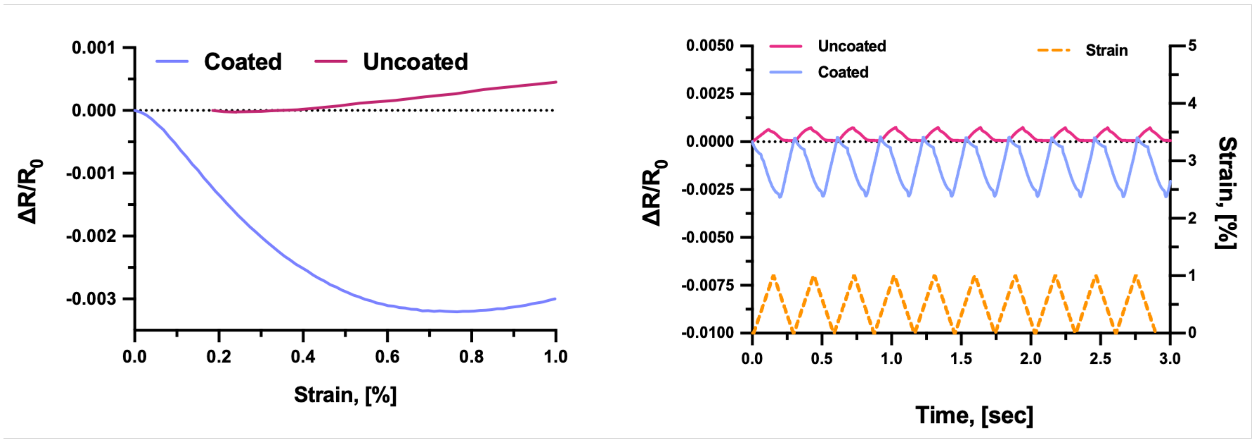

**Figure S7. Sensor signal from pre-strained fibers with tensioned or non-tensioned configuration.** Strain-dependent sensor response of pre-strained fibers under tension (red line) or in loose configuration (blue) in the 0-1% strain range (left), and response under cyclic 1% strain application (right), showing that negative values response can be generated when the fiber is not tensioned at time 0.

Pre-straining the fiber before coating augmented the resistivity as calculated under small (1-2%) strains application (**Fig. S8**), thus indicating that using pre-strained fibers could serve to generate sensitive sensors that are useful to operate on bio-actuators. Coating the fiber reduced the GF, while integrating the fiber into a tissue organoid and keeping it in the cell culture environment for 7 days increased the GF, possibly due to the stiffening occurring in the developing tissue.(3) This observation suggests that integrating the sensor in an embedding matrix with similar Shore hardness would yield high GF, thus maximizing the sensitivity.

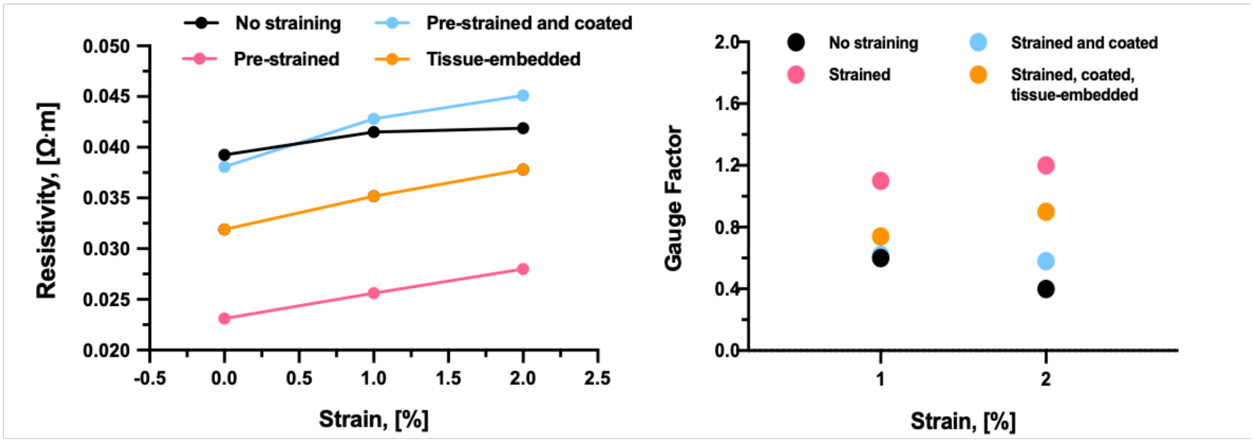

**Figure S8. Variation of resistivity with fiber pre-straining. (A)** Resistivity values were calculated at 0, 1, and 2% strain for non-strained fiber, pre-strained fiber, coated pre-strained fiber and the coated pre-strained fiber that was inserted in a skeletal muscle tissue organoid and remained embedded into it for one week. (**B**) Gauge Factor values were calculated at 1 and 2% strain for non-strained fiber, pre-strained fiber, coated pre-strained fiber, and the coated pre-strained fiber that co-developed together with a skeletal muscle tissue organoid for one week.

### 2. Biocompatibility of the sensor materials

To evaluate the fiber biocompatibility, we used different assays that assess cytotoxicity, and cell adherence. We exposed myoblast cells to different concentrations of CB-polymer composites and for different time ranges (**Fig. S9**). At any incubation time and material dosage, cell viability was above 90%, demonstrating that the fiber is highly biocompatible. Importantly, high cell viability was measured on cells exposed to constituent materials of the coated fibers, suggesting that coating the fiber with silicon did not imply any risk of cytotoxicity. No statistically relevant difference was observed between the cell viability of cells exposed to the constituent materials of the silicon-coated fibers and those exposed to uncoated fibers' materials. Even under the harshest conditions, namely the longest incubation time and highest dosage, the viability only dropped by 10 and 15% for the coated and uncoated fiber, respectively. No statistically relevant difference was observed between the cell viability of cells exposed to the constituent materials of the silicon-coated fibers and those exposed to uncoated fibers' materials.

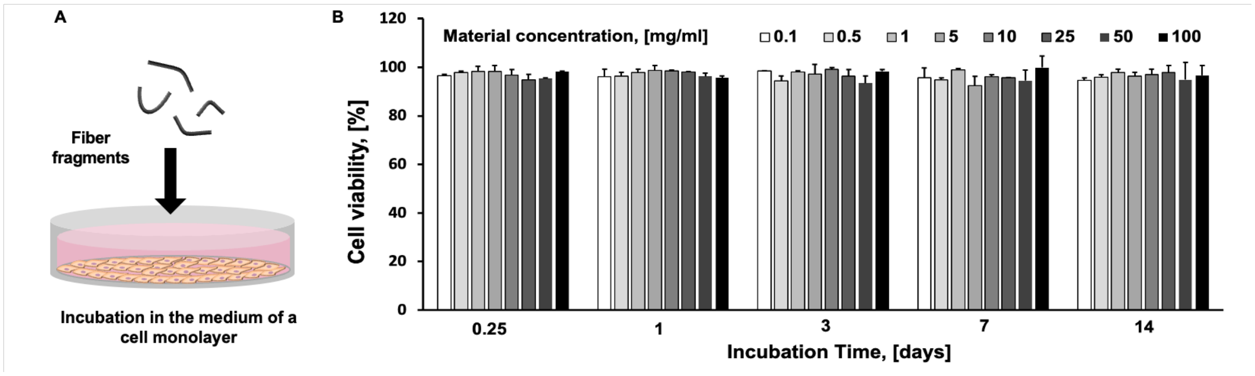

**Figure S9. Biocompatibility of the uncoated fiber on myoblast cells. (A)** Scheme of the cell viability test and **(B)** percentage cell viability measured via a Resazurin assay after exposure to different concentrations of the constituent materials of the uncoated fibers.

Moreover, after 3 h of incubation at intermediate material dosages (10 mg/mL), all cells showed growth trends that were comparable to those of control cells incubated with equivalent volumes of PBS (**Fig. S10**), thus indicating that none of the two fiber types altered the cell proliferation ability.

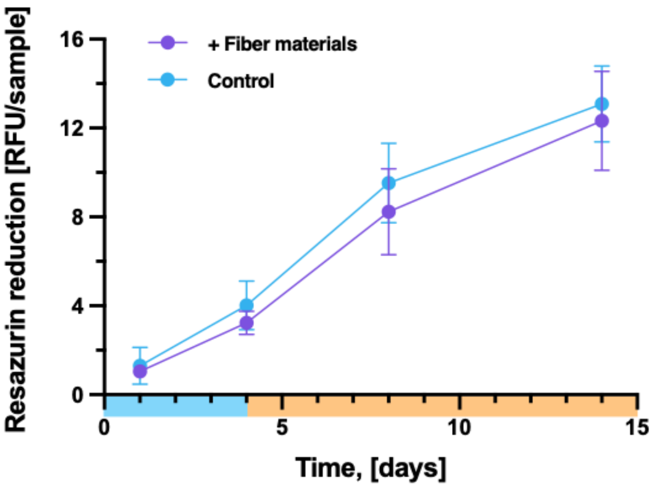

**Figure S10. Myoblast cells proliferation on the uncoated fiber.** Cell viability measured via a Resazurin assay after exposure to 10 mg/mL of the constituent materials of the uncoated fibers.

To complement cell viability information with data concerning cell morphology and adherence, we observed the fibers in Scanning Electron Microscopy, which revealed a dense biofilm formation. Even if some apoptotic bodies were sporadically observed, most cells featured fibroblast-like morphology, cell-cell contact areas, and cellular protrusions for substrate adherence, which confirmed that cell viability and functionality were preserved (**Fig. 3D**). Then, we performed a Live/Dead staining of cells that were seeded and cultured onto intact fibers for a few days. In addition to fibroblasts, we tested cells with electrically excitable membranes that are often involved in bioelectronics, such as skeletal muscle cells and motor neurons. A few hours after seeding, cells adhered to the silicon-coated fibers and displayed high cell viability (> 85%), regardless of their human or murine origin and their culture phenotype (primary cells, hybridomas, or immortalized cell lines) (**Fig. S11**). Myoblast cells on both coated and uncoated fibers displayed high cell viability (> 90%) along one week of culture (1, 3, and 7 days) and a spread morphology, which suggested efficient adhesion to the surface of the fiber (**Fig. S11**).

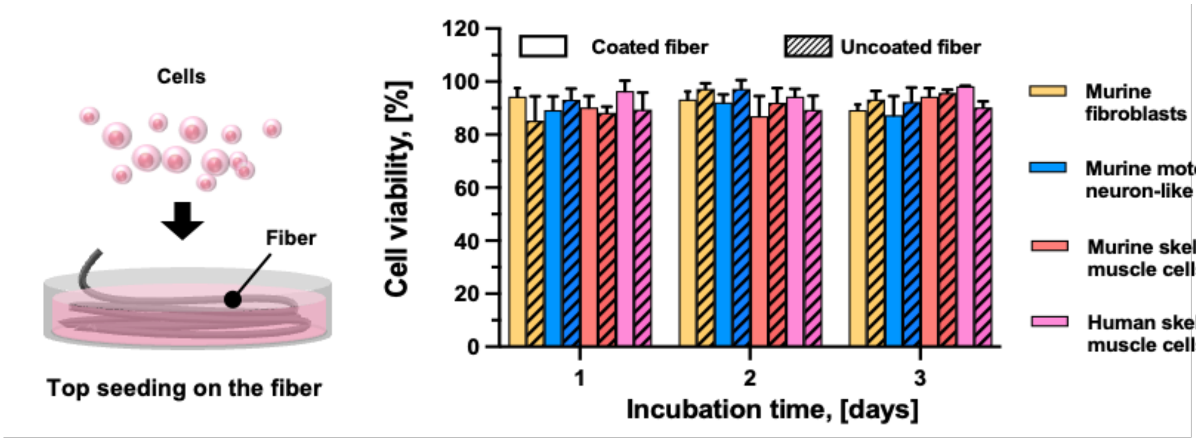

**Figure S11. Biocompatibility of the sensor with different cell types.** Scheme of the incubation assay, in which the cells were let adhere and grow on the fiber (left). Cell viability measured via Live/Dead staining on cells adhering onto intact fibers, such as murine fibroblasts (NIH 3T3), skeletal muscle cells (C2C12) and motor neuron-like cells (NSC-34), and human primary skeletal muscle cells (right).

### 3. Assembly into 3D tissue culture models

To understand if the sensor could be integrated with a developing 3D tissue model, we embedded the coated fiber into a myoblast-laden hydrogel construct (1×0.5×0.5 cm) during the biofabrication process (**Fig. S12**), and analyzed the cell proliferation and differentiation ability, as well as the skeletal muscle tissue maturation process. The hydrogel gelated around the fiber, and tissue formed around it without losing integrity. **Fig. S12** shows the procedure to fabricate the tissue-integrated fiber construct: the fiber was inserted in a custom-made mold of rectangular shape with a break at the walls to let the fiber slip down. To maintain a pre-tensioned state, the fiber was tended at its extremities during the gel formation process. A Collagen-Matrigel mixture seeded with $100\times10^6$ cells was slowly cast around the fiber and let gelate around it in the incubator. The constructs were carefully removed from the mold and sat in cell culture wells with appropriate conditions for skeletal muscle tissue development. Immediately after gelation, the constructs appeared to have an inhomogeneous matrix containing air bubbles and cells (appearing as roundish bodies, as not yet adherent to the matrix surface).

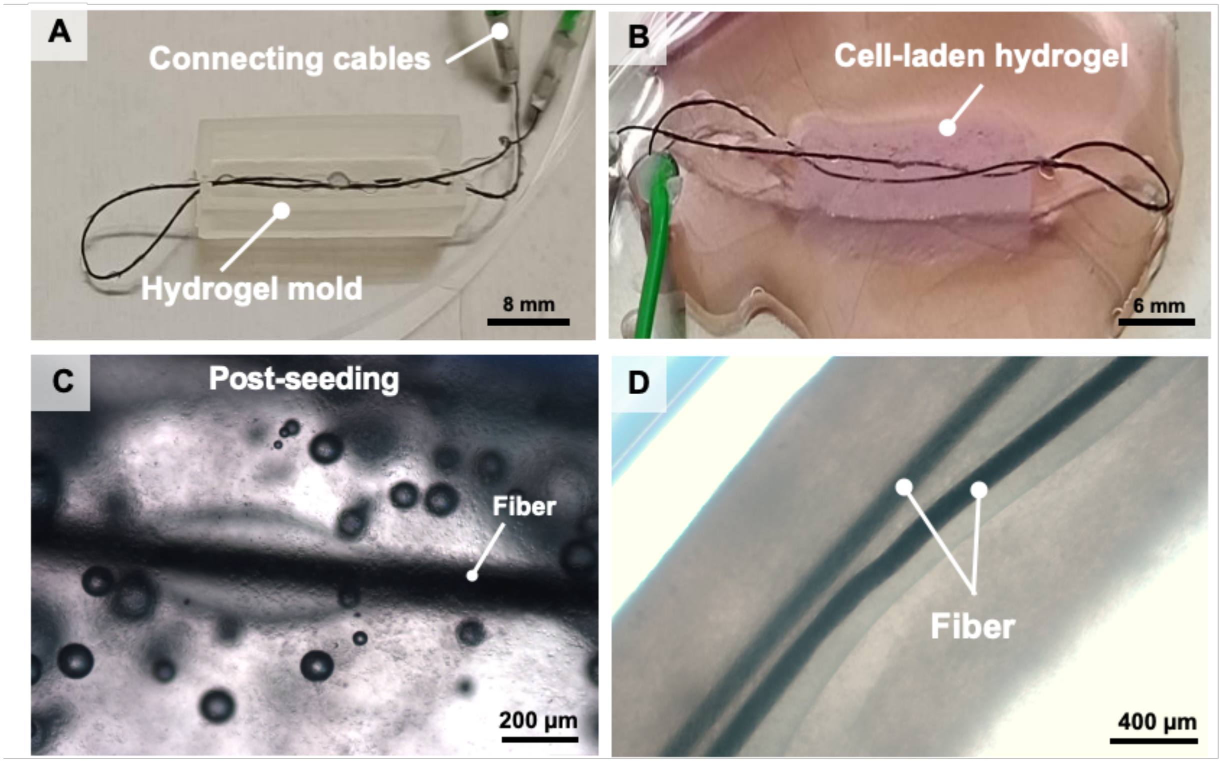

**Figure S12. Fabrication of the sensorized skeletal muscle tissue.** Steps to fabricate tissue-integrated fiber: optical picture of the looped fiber that is inserted within the mold before pouring the cell-laden hydrogel (**A**) and fiber-integrated construct immediately after demolding (**B**). Microscopic imaging of tissue-integrated coated fibers immediately after molding (**C**) and 15 days post-biofabrication (**D**). After seeding, microscopy reveals details of the coating and cells with round shapes. After tissue development, fibers are visible within the tissue matrix.

Cells were homogeneously distributed all over the construct volume (**Fig. S13**).

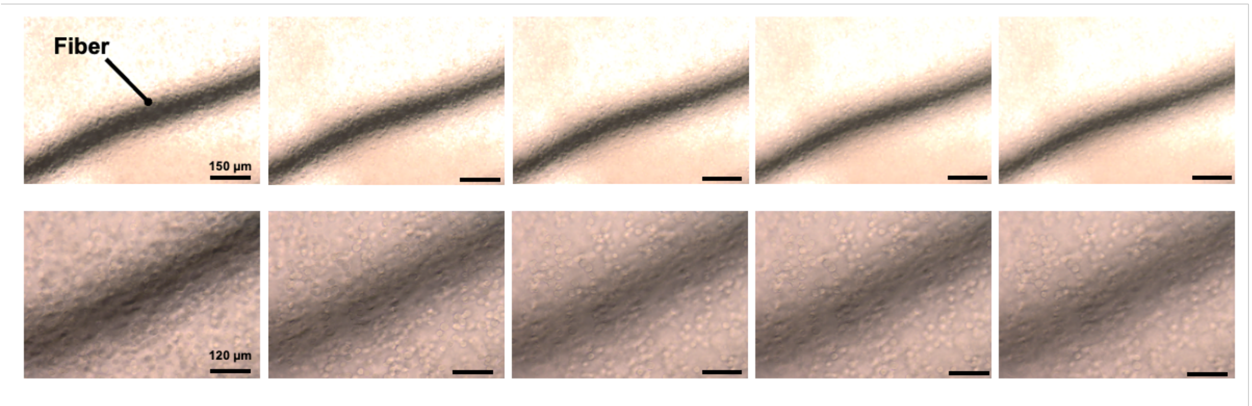

**Figure S13. Cell seeding within fiber-integrated constructs.** Optical microscopy of a fiber-integrated construct immediately after fabrication, showing cells homogeneously distributed at different depths around the fiber (increasingly superficial layers of tissue are shown from left to right). Magnification: 10 and 20× (top and bottom, respectively).

Despite tissue condensation and remodeling, the fiber remained well-positioned for several days in culture, and viable cells were observed via light microscopy at various distances from the fiber, displaying fibroblast-like and elongated morphology after 1 and 7 days of culture, respectively (**Fig. S14**).

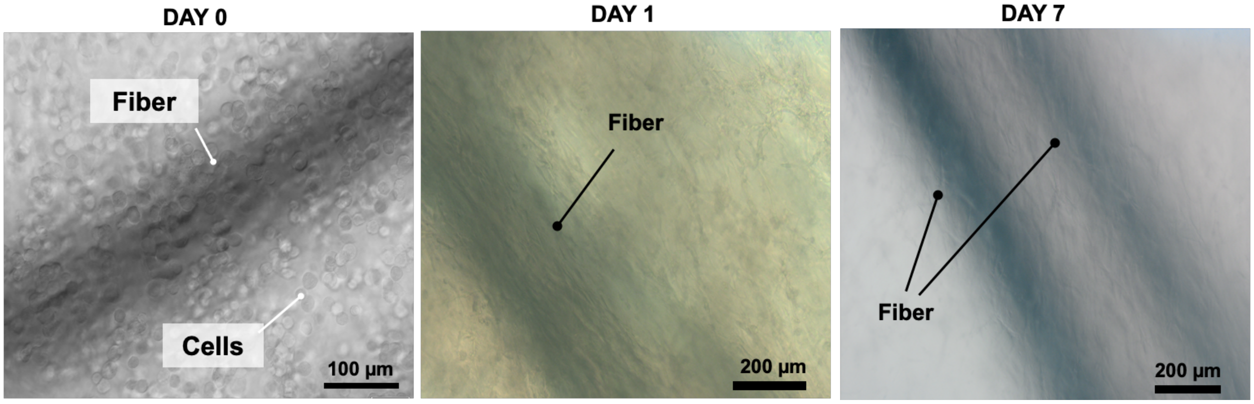

**Figure S14. Fiber's integration and positioning into cell-laden hydrogel.** Optical microscopy of fiber-integrated constructs showing cells dispersed in the hydrogel matrix immediately after hydrogel molding and at days 1 and 7 after fabrication. Immediately after construct seeding, cells displayed a round shape, while after 1 and 7 days of tissue development cells displayed an adherent and elongated morphology, indicating cell viability, adherence, and differentiation.

To gather deeper insights into skeletal muscle tissue maturation, we analyzed the culture media via ELISA to quantify the production of myogenic function regulators, including two ligands for the cytokine receptors (IL-4, IL-6), and one member of the TGF-β family (Myostatin) (**Fig. S15**). All three conditions displayed an increasing cumulative release of IL-6 and IL-4, which promote the satellite cells' activation during early myogenic differentiation, and the second-stage fusion in later differentiation stages, respectively. In contrast, as expected for a negative regulator of proliferation and inhibitor of satellite cells' differentiation, a progressive decrease in Myostatin production was observed in all constructs. This cytokine production profile suggests that cells differentiated into skeletal muscle tissue regardless of the fiber inside the construct.

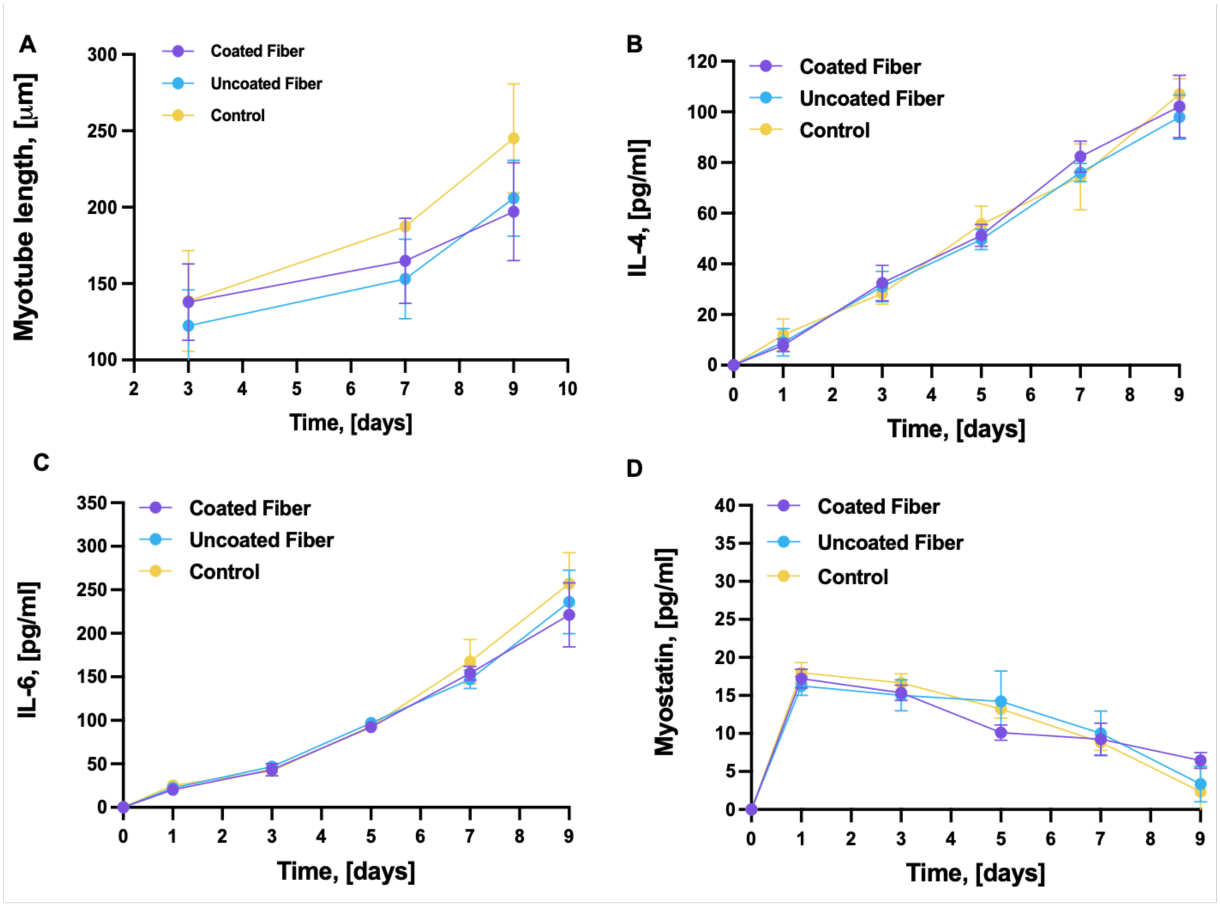

**Figure S15. Skeletal muscle tissue differentiation and secretome analysis.** (**A**) Myotube length as calculated from f-actin staining on confocal imaging (n = 3). Secretome analysis via quantification of cytokines in the culture media: IL-4 (**B**), IL-6 (**C**), and Myostatin (**D**). In all assays, $n = 3$.

### 4. Interfacing with living tissue

To study the durability of the tissue-fiber interface, we fabricated a skeletal muscle tissue organoid and inserted the fiber in the construct at day 1 post-fabrication, thus in the early stage of tissue development. At the end of the maturation process (day 11), a homogeneous tissue matrix was observed, in which elongated cells and myofibers were visible (**Fig. S16**). Importantly, despite the dramatic tissue condensation effects that occur during tissue development, the fiber could retain its positioning within embedding organoids. The matured tissue tightly adhered to the fiber, forming a cohesive interface between the sensor and organoid (**Fig. S16 B-C**).

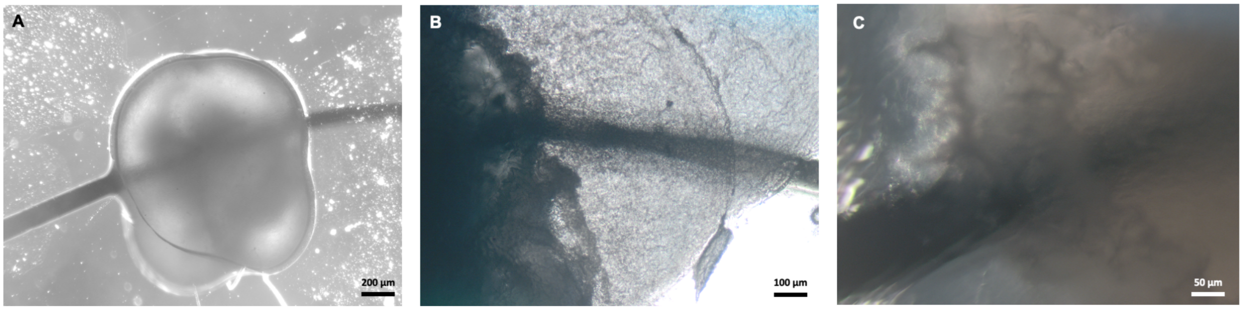

**Figure S16. Sensor integration during tissue maturation.** Optical microscopy of a fiber-integrated construct at the end of the maturation protocol for engineered skeletal muscle tissue, showing (**A**) the fiber included in an organoid that retains its positioning after tissue condensation and remodeling; (**B**) the area where the fiber penetrates the tissue; (**C**) the cohesive tissue-sensor interface on the organoid's surface.

The interface between fiber and matrix in the absence of cells was loose, suggesting that the co-developed living tissue plays an active role in stabilizing the interface (**Fig. S17**).

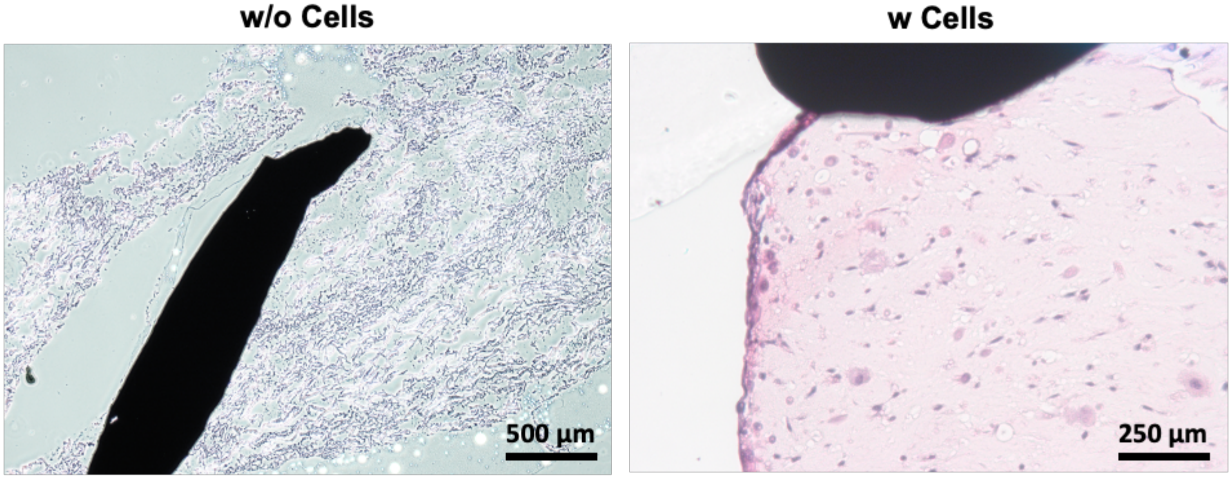

**Figure S17. Sensor-tissue interface formation.** H&E histological staining of fiber-matrix constructs, obtained by gelating a collagen hydrogel around the fiber in the absence (left) or presence (right) of cells. When cells are missing, the fiber has poor adherence to the hydrogel matrix and lacunae are visible on the interface.

Finally, we explored the sensor integration following the tissue development, showing that the fiber can be applied to the formed tissue by gluing it with a biomedical adhesive or sewing it into the tissue (**Fig. S18**).

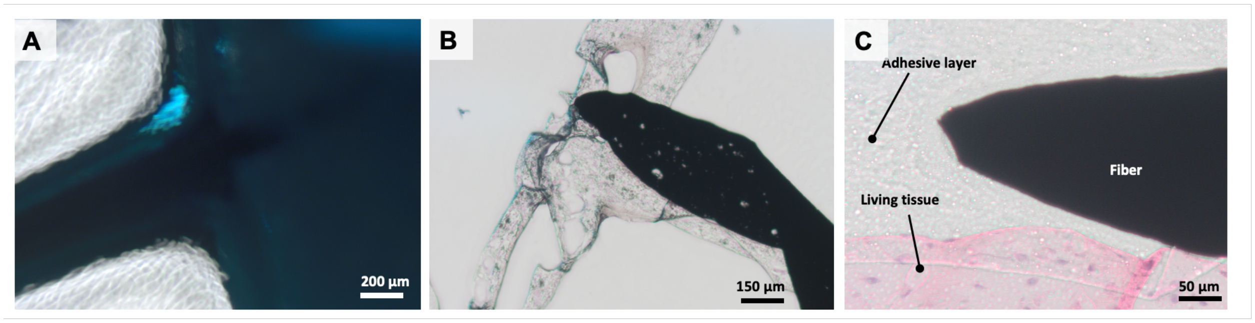

**Figure S18. Alternative tissue-fiber integration approaches.** Optical picture of the fiber glued to a skeletal muscle tissue organoid, in which the glue provides a blue color to the construct as observed under light microscopy (**A**). H&E histological staining showing the blue-colored glue adhering around the fiber (**B**) and the glue-based-intermediate layer that participates in the bio-interface generation (**C**).

### 5. Operation in soft matrices

We aimed to understand how the sensor would operate when exposed to biologically relevant environments. Therefore, we first tested sensor reliability when integrated into soft matrices. We observed that under no mechanical loading, the fiber could efficiently adhere to the irregular surface of hydrogel blocks, establishing tight contact with these materials (**Fig. S19**).

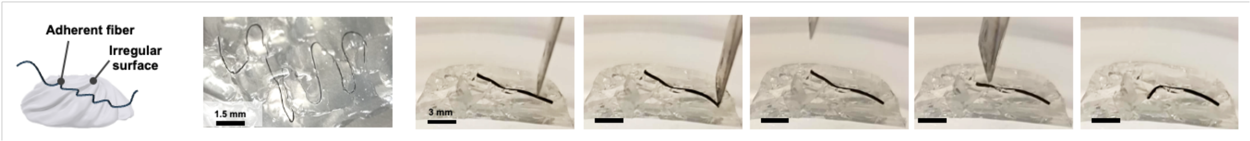

**Figure S19. Flexibility and adhesive properties of the fiber**. Optical pictures of the fiber adhering to irregular hydrogel surfaces under gentle touch with a spatula.

To understand how the embedding of the fiber in a soft matrix affected the sensor response under realistic loading conditions (tension, compression), we studied the fiber's integration and functionality within soft matrices of the appropriate size used as tissue-mimicking phantoms. By measuring the solidity of soft tissue phantoms with a durometer, we found that 1.5% agar-based phantoms have a similar solidity to the skeletal muscle tissue that is engineered from a collagen and matrigel hydrogel mixture and cultured for several days in the lab to form matured tissue (≈ 7.5 Shore A). In contrast, the hydrogel mixture without cells displayed lower Shore hardness values (≈3A) that did not significantly differ after cell addition (≈3.5A) (**Fig. S20**). When embedded into 7.5A phantoms, the fiber compliantly followed the structural deformation that resulted from compressing the soft matrices in different ways (**Fig. S20**).

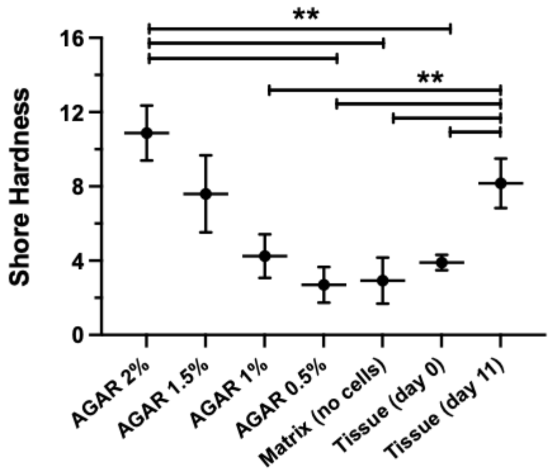

**Figure S20. Softness of matrices used for fiber integration.** Shore hardness measured for soft matrices, including hydrogels for skeletal muscle tissue engineering, agar-based tissue phantoms, and fully matured engineered muscle tissue.

Two different configurations were examined for the position of the fiber within the phantoms; in one case the fiber was kept straight, while in the other case, the fiber was looped to form a serpentine arrangement (**Fig. 4A** and **C**). We observed that the fiber compliantly bent with the tissue phantom (**Fig. 4A** and **C**), without causing damage to the surrounding matrix. When investigating whether applying the mechanical stimuli to the matrix could be coherently perceived by the embedded fibers, we observed that embedding the fiber in a straight configuration within non-cell populated matrices resulted in unreliable sensor behavior, thus suggesting the interface between the matrix and the sensor was inefficiently stabilized. It likely resulted in the slippering of the fiber within the agar phantom. As such, we mechanically stabilized the integration of the fiber by arranging it in a pre-tensioned serpentine design that could remain embedded in the gel block during applied mechanical stress (**Fig. 4C**).

By embedding the coated fiber in different matrices and applying strain stress (≈70% of the initial fiber length) to the fiber, we observed that the relative resistance values varied in the 0.045 - 0.065 range for the different tested materials (**Fig. 4**). In particular, the values of relative resistance variation measured for fibers inserted in engineered skeletal muscle tissue and agar phantoms with a similar solidity were comparable (0.055 ± 0.009 and 0.047 ± 0.01, respectively) and close to values of fibers that were not embedded in any matrix (0.048 ± 0.002). No significant difference was found between the sensor responses of fibers embedded in tissue constructs at different matrix maturation points (days 1 and 11) (**Fig. 4B**). Thus, we conclude that embedding the fiber in soft matrices and culturing in media did not significantly alter the fiber responsivity to the applied strain stimulus.

To investigate if applying the mechanical stimuli to the soft matrix could be coherently perceived by the embedded sensor fiber, we stretched and compressed the agar-based tissue phantoms by ≈30% of their length and measured the relative resistance variations in the fibers aligned along the direction of the applied mechanical force (**Fig. S21A** and **B**). By repeating the phantom compression over time, we observed that the relative resistance increased without any defined strain-dependency trend, thus confirming that the sensor could be more reliable in detecting strain stresses than the compression ones. Despite the presence of significant signal relaxation, possibly due to the viscoelasticity of the matrix, it was possible to distinguish the time points of load application and load removal. The strain response of the embedded fiber was lower than what was observed in the characterization data reported in **Fig. 2F**. In addition, when applying the 100g load for the first time, the sensor response matched the profile observed at the test without the weight. However, when loading the weight afterward (200g), the sensor response was not the expected one, and recognizing the time points of load application and removal was difficult. Consequently, we assumed that upon the first load, a dislocation of the fiber inside the matrix occurred resulting in an inaccurate detection of the second load. This suggested that the stimulus could not be efficiently transferred from the matrix to the fiber, likely due to the slippering effect of the fiber within the phantom after the deformation caused by the first load. We subjected the fiber to strains derived by applying 0.98 and 1.96 N forces generated from 100 and 200g weights applied on the block surface and corresponded to fiber elongations of ≈3% and 5% (**Fig. S21C)**. The sensor response did not recover its background values, which indicates that large fiber deformations could result in the plastic behavior of the materials. Of notice, when embedding the fiber inside soft matrices, the behavior of the sensor presented certain peculiar features. For example, while a necking effect was observed when the uncoated fiber was brought to the yield point, this effect was not reported for the coated fiber. This could suggest that in the coated fiber, a slipping effect could occur starting at low strains.

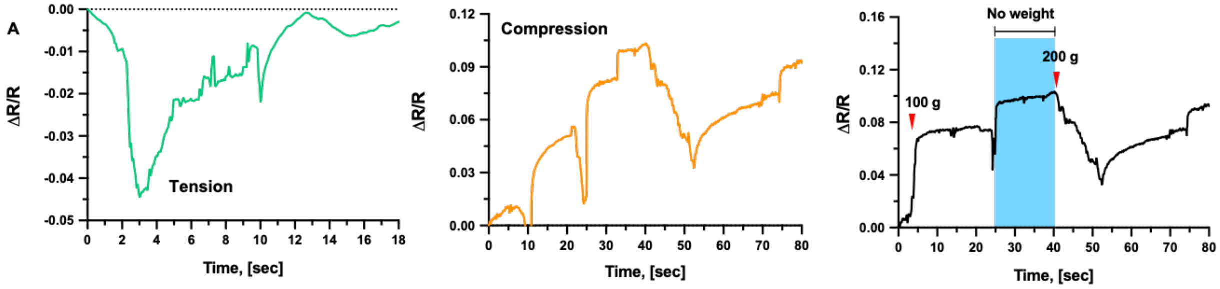

**Figure S21. Force transmission across soft matrices and sensor performance.** Time-dependent variation of the relative resistance of the gel-integrated fiber under compressive (**A**) and tensile (**B**) stimuli, and under consecutive application of two compression stimuli (two weights of 100 and 200 g, indicated with red arrows) (**C**). Blue areas indicate a time range where no weights were applied, potentially reflecting relaxation processes occurring in the phantom construct.

Compressing and tensioning of the phantom block caused a sensor response from the serpentine fiber, which provided negative and positive relative resistance responses, respectively (**Fig. 4D** and **E**). This response did not match either the behavior of the fiber tested in the air (**Fig. 2**) or the one of the fibers embedded in the soft phantom and arranged with a straight design. The different behavior could be ascribed to the serpentine design, in which the orientation of the sensor materials to the applied stress is diversified and causes a complex resistivity response. However, through many convolutions, this design avoided the slipping of the fiber within the matrix, thus serving to better anchor the sensor. While data collected from the compressed phantom were noisy, those generated under the tensile stimuli were clear (**Fig. 4D** and **E**). We associated this difference with the fiber arrangement and the difference in the loading. Under tension, no noise was observed (**Fig. 4**), thus matching the quality of the sensor response observed during the tensile testing (**Fig. 2**). Repeating the phantom tensioning resulted in a coherent sensor response, as the frequency of the signal peak pattern matched the one of the applied stimuli paradigm (**Fig. 4F**).

### 6. Operation in the cell culture environment

To understand if the sensor could operate within biological environments, we then assessed the sensing performance of the fiber after long-term exposure to conditions required for cell culture that simulate *in vivo* environments. After sitting in cell culture media at 37°C for one week, the strain-dependent stress and relative resistance response of the fibers differed from the ones observed in the basic characterization conducted in the air (**Fig. 2, S22**). The stress responses of sensors that were exposed to the cell culture environment had positive but lower values (≈ 2-2.5 MPa vs. 4-10 MPa range at 100% strain) and different curve shapes (**Fig. S22**). A different behavior between coated and uncoated fibers was also observed. As shown in **Fig. S22A**, after exposure to culture media, the fibers' stress responses had different curve shapes with positive but lower values (≈ 3 MPa vs. 4.5 MPa range at 200% strain for uncoated and coated fibers, respectively) as compared to those of the basic characterization in air. Submerging the fibers in the medium likely resulted in weighting the fiber down, thus having a pre-loading effect on the stress profile during the tensile test.

After incubation in cell culture media, the relative resistance response of the uncoated fibers switched from positive values with a strain-dependent increasing profile (**Fig. 2**) to a non-monotonic response (**Fig. S22C**), suggesting that, in a biological environment, possible deterioration of the fiber structure and composition occurred due to the absence of the coating. Even if the behavior of the coated fiber also differed after the long exposure to the cell culture environment, it displayed a negative response with a monotonic trend (**Fig. 2F, left**), which could serve to generate a useful sensor response in a defined strain range (*i.e.*, 0-30%).

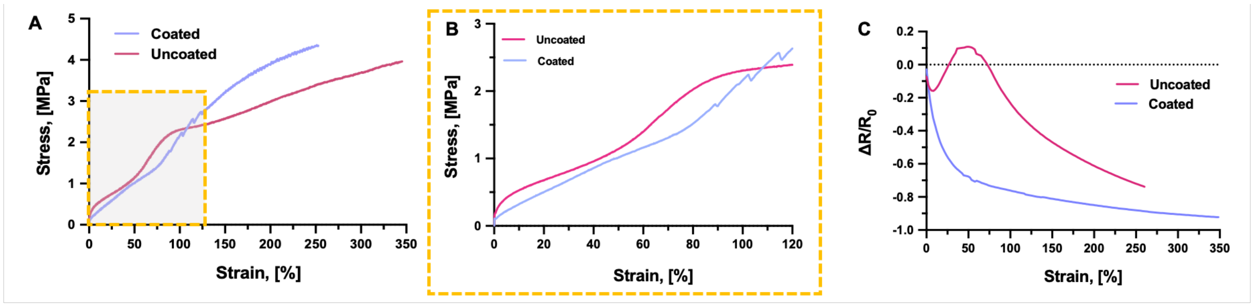

**Figure S22. Sensor performance after exposure to cell culture environment over a wide strain range.** Stress-dependent variation of the stress on the 0-350% (**A**) and 0-120% (**B**) strain ranges, and (**C**) the relative resistance of coated and uncoated fibers after 7 days of incubation in cell culture media at 37°C.

We studied the fibers' dynamic responses to repeated 50% strains (**Fig. S23**). The coherent behavior between stress and resistance response observed in the dynamical tests proves that the sensor response is reliable. after exposure to conditions that are relevant to integration with engineered tissue and other biomedical applications, such as repetitive stresses.

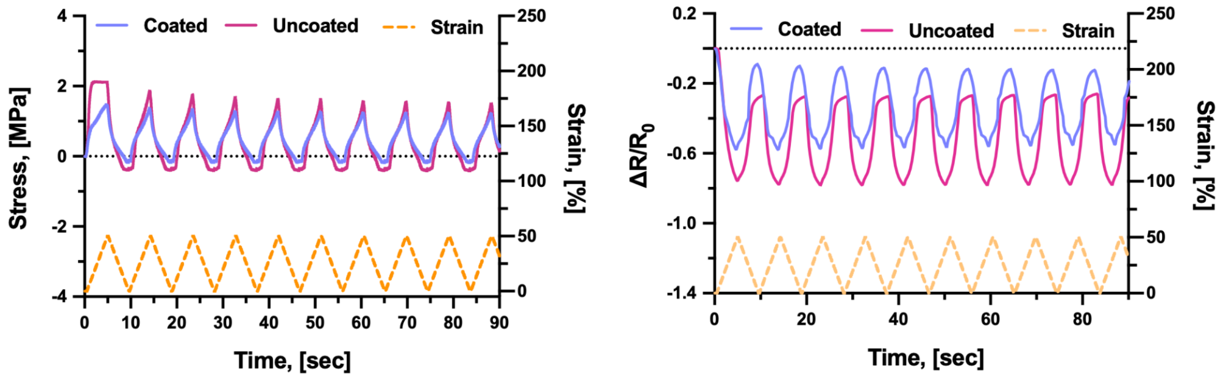

**Figure S23. Sensor dynamic response after exposure to cell culture environment.** Time-dependent variation of the stress (**left**) and the relative resistance (**right**) of coated and uncoated fibers under strain stress (applied at 50%) after 7 days of incubation in cell culture media at 37°C.

In addition to the physical conditions of the cell culture environment, the cell activity (e.g., nutrient consumption, metabolite production, etc.) was investigated in its effects on the sensor operation ability over time. To assess the sensing performance of the fiber after long-term exposure to biological environments accounting for cell activity, the fiber was mounted on a custom-made setup that enabled the tensioning of the fiber in a dish with myoblasts cells growing in monolayer (**Fig. S24A**), and then tested for strain-response after 7 days of exposure to cell culture media and products of cell bioactivity (**Fig. S24B** and **C**). With straining, the relative resistance of the coated fiber increased with a trend that resembled the strain response measured in air (**Fig. S24D**) but with lower values (0.44 vs. 2.0 Rel. Res. at 20% strain for the cell culture and air measurements, respectively). The response of the uncoated fiber presented incongruent data points that suggested operation unreliability in the cell culture environment. In contrast, as the strain-dependent variation of the relative resistance of the coated fiber matched the strain response measured in air (**Fig. 2F**), when coated fibers were exposed to an active cell culture environment in which the medium's composition was affected by the cell bioactivity, they retain their functionality. As the uncoated fibers displayed operation unreliability, we concluded that the coating contributed to preserving the fiber and its function. The sensor response resembled the response observed during the test of the phantoms (**Fig. 3**). Upon static strain application, the resistance values reached their plateau in a subsecond time range, exhibiting a relaxation behavior of the electrical signal, a well-known effect attributed to the viscoelasticity of the elastomeric matrix material (**Fig. S24C**) (3,4). When detecting the rapid mechanostimulation response induced by a contracting bio-actuator, the sensor signal relation is not expected to majorly impact the sensor's response and a sensor response more similar to the one characterized during the dynamic test (**Fig. 2G**) would likely occur. In contrast, a sensor signal relaxation is expected when the bio-actuator is resting (e.g., in the absence of electrical stimulation) (**Fig. S24**).

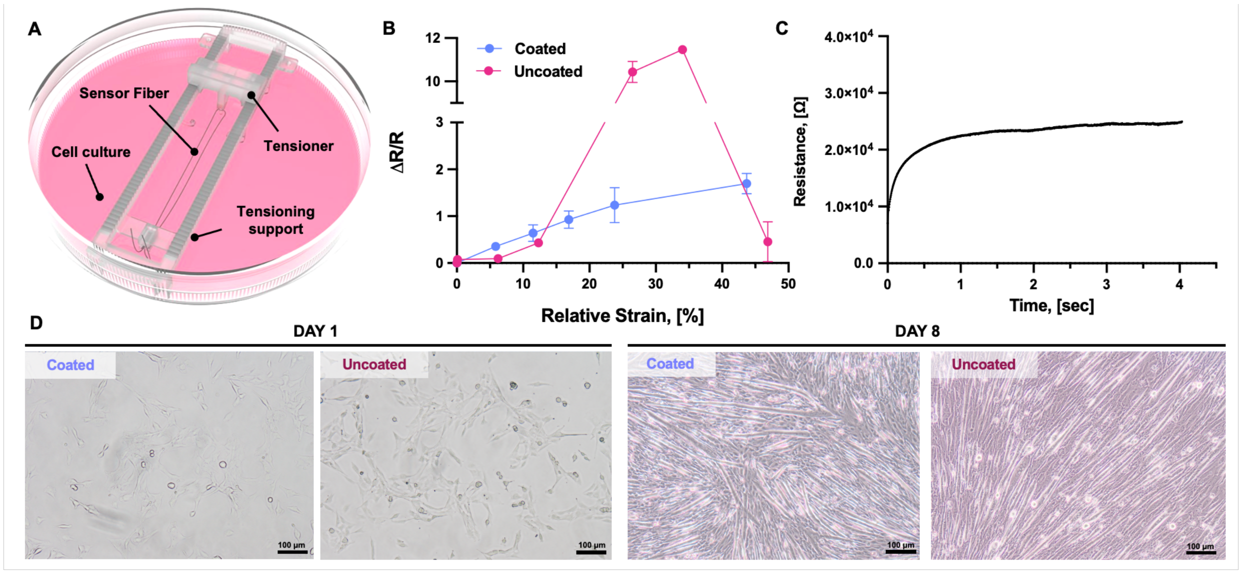

**Figure S24. Sensor performance and effect on 2D cell culture.** (**A**) Setup for fiber tensioning within liquid media of a cell culture dish containing a monolayer of myoblasts. (**B**) The relative resistance change of coated and uncoated fibers under strain stress after 7 days of exposure to a cell culture environment. (**C**) Example curve of resistance measurement of coated and uncoated fiber at day 7, under relative strain respectively. (**D**) Optical microscopy shows viable cells growing on day 1 after resistance measurement and myotube formation on day 8 after the second resistance measurement.

Finally, when exposed to applied electrical fields in a cell culture medium, the uncoated fiber's response presented an electrical drift of 6% between the different pulse cycles. In contrast, the relative resistance of the coated fibers was mostly unaffected by the electrical pulses, and their electrical drift was 2% only. The coating successfully shielded the fiber from the applied voltage, thus we assumed that any changes in the signal would be caused by the piezoresistive properties of the fiber in the bionic construct.

### 7. Amplification of the sensor signal from the bio-actuator's micromotions

As the limited contraction motion and forces involved in the bio-actuators' response could generate noisy signals, we defined an appropriate processing approach to render minimal sensor signals useful in effectively controlling bio-actuation dynamics (**Movie S1**). After filtering, the processed sensor signal was used to instruct a posteriori control system to stop the output of electrical stimuli in response to the contracted states of the stimulated muscle (**Fig. S25A**).

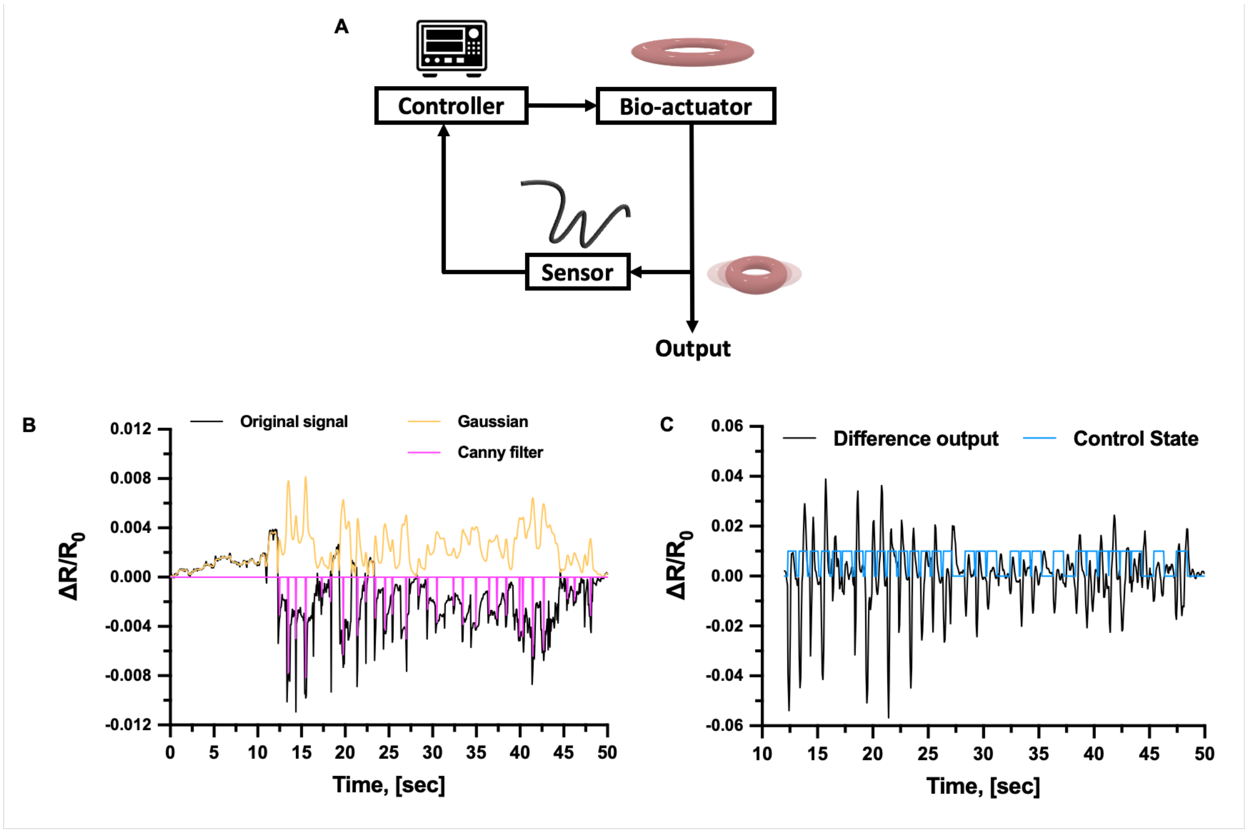

**Figure S25. Sensor signal data optimization.** (**A**) Scheme of the control circuit around the bio-actuator's activation. (**B**) Original and processed signal from sensor response of the bio-actuator plotted with the control signal under 1 Hz stimulatory pattern. (**C**) Output signal difference plotted with the control state.

The original sensor response shown in **Fig. S25B** (black line) was filtered through a series of Gaussian filters (orange line), and a Canny Edge detector was applied to the resulting signal to determine points of maximum contraction (shown in pink) on the original signal (**Fig. S26**).

**Algorithm 1: Open loop analysis**

$R_{all}$ ← Array with all obtained measurements
$M_{all}$ ← Array with all found minima
Apply three consecutive Gauss filter $\mathcal{N}(\mu, 1)$ over $R_{all}$
**procedure** Canny Edge detector
  **repeat**
    Sample data from with $window = 7$
    **if** minima in sample:
      $M_{all} = M_{all} \cup M_{new}$
    **end if**
    Move window to 7 steps
  **until** all data sampled
  $M_{all} = \frac{M_{all} - M_{all_{min}}}{M_{all_{max}} - M_{all_{min}}}$
**end procedure**
Find frequency of minima
Compare frequency to original control signal

**Figure S26. Open loop analysis algorithm.** Pseudocode of algorithm used to process raw resistivity measurements from muscle contractions under stimulation.

The approach showed clear minimum identifications with high correspondence (approximately 78.5%) with the originally applied control signal. The missed points were attributed to a high presence of noise in the measurement which resulted in ambiguous dense minima sections. We calculated a control output signal a posteriori by subtracting the current filtered measurement with the measurement obtained five-time steps prior (in black, in **Fig. 25C**). The difference output signal was used to determine the behavior of the controller. The controller was configured to trigger a simulated signal at 1 Hz and analyze the rate of change of relative resistivity in the fiber to determine the shut-off point on each trigger cycle. The designed controller successfully realized the expected cut-off action at the beginning of contraction that matches the observed behavior in the data analysis with 100% efficacy (maintaining a correspondence match of 78.5% to the ground truth). The missed expected behavior of the controller was due to the presence of noise, which disturbed the applied thresholding (-0.006). Thus, as the pattern of the controller response (in blue, in **Fig. 25C**) matched the profile of the processed sensor signal, we conclude that after appropriate processing, the sensor signal would be useful in effectively controlling bio-actuation dynamics.

### 8. Closed-loop control of the bio-actuator

To understand whether the sensor could be used to control the behavior of a bio-actuator, we built a platform in which the sensor was integrated into the maturing skeletal muscle tissue. After a 24-hour growth period, a ring-shaped bio-actuator was looped at its two extremities with two sensing fibers that were wrapped around pillars (**Fig. S27**). The ring was subsequently tensioned and left in culture for the required tissue maturation time.

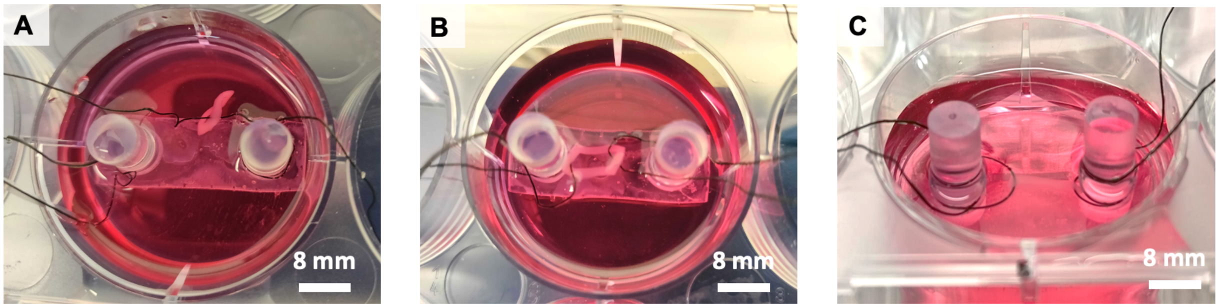

**Figure S27. Mounting procedure of the sensorized bio-actuator platform.** Optical pictures of the assembly of the sensorized bio-actuator's platform, starting from inserting the first (**A**) and second (**B**) fiber in the ring-shaped actuator. (**C**) By tensioning the fiber, the bio-actuator was subjected to applied tension.

By applying electrical fields to cell culture media, the bio-actuator's contractile response was obtained (**Movie S2**) and used to determine the threshold values to apply to the control system. The actual controller behavior (**Fig. 6**) followed the expected behavior, operating an on/off switch action with sull reliability (100% of expected controller responses) (**Fig**. **S28** and **Movie S3**).

**Algorithm 2: Closed loop controller**

$R_{all}$ ← Array with all measurements
$T_{rate}$ ← Trigger rate $[triggers/s]$
**repeat**
  **if** $T_{rate}$ % $time_{current} = 0$
    *Output ON*
  **end if**
  Get resistivity measurement $R_{rel_t}$
  $R_{all} = R_{all} \cup R_{rel\_t}$
  Apply Gauss filter $\mathcal{N}(\mu, 1)$ over $R_{all}$
  Apply weighted median filter ($[1, 3, 5, 3, 1]$) on $R_{all}[t-5]$ $to$ $R_{all}[t]$
  $\Delta R_{rel_t} = R_{rel_t} - R_{rel_{t-5}}$
  **if** $\Delta R_{rel_t} > Threshold$
    *Output OFF*
  **end if**
**until** controller off

**Figure S28. Closed loop controller pseudocode.** Steps taken to realize the closed loop controller.

When no stimulation was applied, the controller's system could not operate as no sensor response was detected (**Fig. S29** and **Movie S4**). The baseline signal did not significantly differ from the open and closed-loop systems (**Movie S2** and **S3)**.

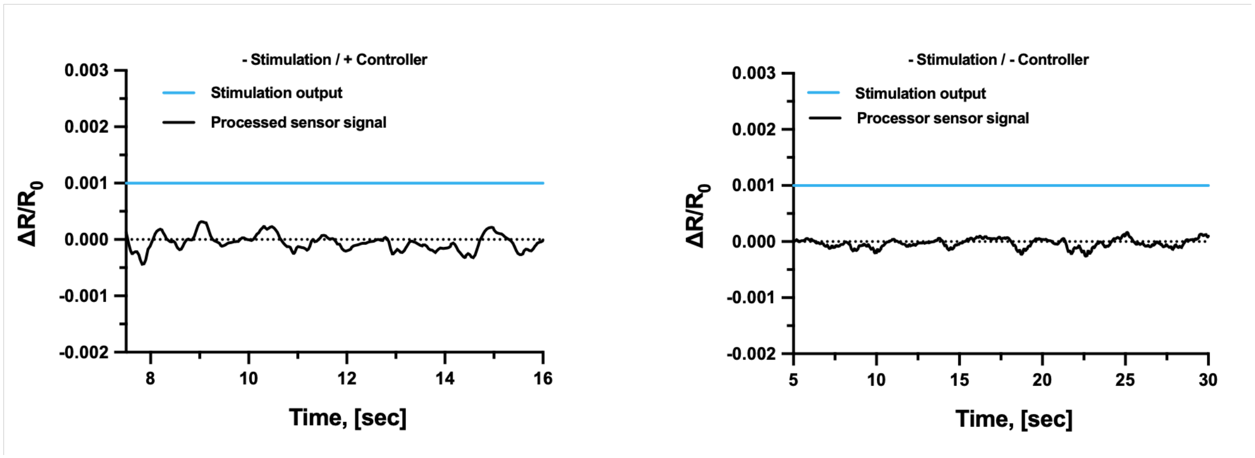

**Figure S29. Baseline signal and controller behavior.** Processed sensor signal (black line) and real-time behavior controller (blue line) in closed and open-loop systems (left and right, respectively). No sensor response was detected and the control system did not operate.

When applying electrical stimulation, the system successfully performed with different triggers (1, 3, and 5 secs) and different thresholding (**Fig. 6H, J, K**, and **Fig. S30**), matching bio-actuator's response patterns with evident or less evident rhythmic signals (**Fig. 7**).

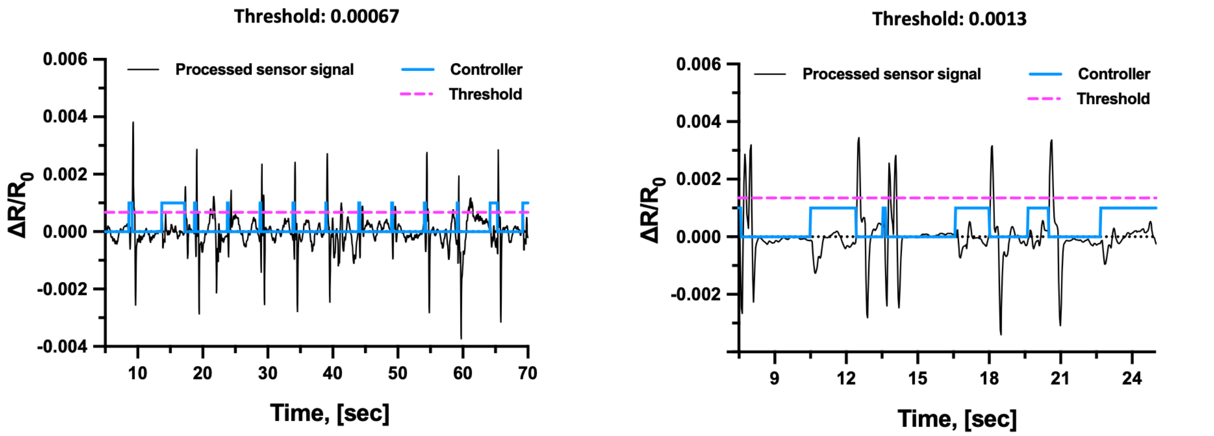

**Figure S30. Closed-loop system with different thresholds.** Processed sensor signal (black line) and real-time behavior controller (blue line) in closed-loop actuation performed with thresholds set at 0.00067 and 0.0013 (left and right, respectively). The threshold value is shown (dashed pink line). Sensor response of the bio-actuator and controller behavior under triggers set at 5 secs.

### Extended discussion

Bio-hybrid robotics has emerged as a frontier area of robotics looking at soft, renewable, and intelligent materials for compliant motion realization (5). In particular, bio-actuators participate in the soft robotics' mission of developing artificial muscles that match or mimic the performance of native muscle tissue (5,6). Bio-actuators are expected to overcome the inability of synthetic materials to reproduce the hierarchical architecture of muscle tissue, thus matching its functional complexity and force output (7). Nevertheless, one of the major challenges for future bio-hybrid robotics concerns the realization of motion control autonomy, which starts from informing the system of its dynamic state (8). With our work, we took the first step towards intelligent autonomous motion by demonstrating closed-loop control of bio-actuator contraction, achieved by the integration of a soft and biocompatible fiber-shaped sensor into the bio-actuator. studying the sensorization of the bio-actuators with emphasis on the biocompatible composition of bioelectronics, its ability to integrate into soft tissues, and the operativity in cell culture and cell stimulation environments.

Our first main achievement concerned the development of tissue-integrated mechanical sensors. Mechanical sensors are garnering relevance in biomedical engineering due to their ability to detect and measure mechanical forces and deformations in biological systems, biomedical models, and biometrical devices (9–11). For instance, mechanical sensors for continuous vital signal monitoring and detection of altered physiological status have been developed, which include wearable sensors that can noninvasively and dynamically collect real-time biomechanical information with a high degree of conformity (10,12). Mechanical sensor technologies with prospective applications for smart, implantable devices have also been proposed (13,14). Direct contact or implantable mechanical biosensors can sense physiological parameters and quantify tissue response, thus allowing one to design personalized treatment plans based on the specific patient's needs. However, for use in bioengineering applications, component materials, designs, and manufacturing approaches of mechanical sensors have to be optimized to improve the performance of the sensors (*i.e.*, sensitivity, durability) within biological environments that feature complex architectures, soft materials, liquid milieu, and activity of living systems (*i.e.*, cells) (15,16). Developing sensors that are biocompatible, reliable, and minimally invasive poses a significant challenge, which limits their application to natural or engineered biological systems (17,18). However, if we could efficiently incorporate sensing units into fragile bio-systems and combine them with simple readout methods, embedded mechanical sensing would allow us to monitor the structural stresses and control motion, as well as understand tissue repair and formation processes (19,20).

Recent advances in micro- and nanotechnology have enabled sensors with flexible, miniaturized, and less invasive designs, thus creating new opportunities for direct contact and implantable sensor technologies (21). Various mechanical sensor types have been used for biomedical engineering applications including piezoelectric, capacitive, and piezoresistive sensors. Due to their versatile strain-sensing abilities, easy readout and low energy requirements, piezoresistive materials have been widely investigated for robotics, biomedicine, and other applications (*e.g.*, for biometrics and prosthetics), and enable wearable electronic devices, electronic skin, and flexible sensors, and implantable interfaces (22,23). In particular, due to their mechanical properties and low production costs, piezoresistive elastomer-based composites are attractive sensor materials that can be integrated within soft and flexible systems. These composites are obtained by integrating conductive filler materials in an elastomer matrix with the desired processability and Shore hardness, and their piezoresistive behavior is caused by the electron tunneling effect, electron hopping and the separation of conductive filler elements within the percolating network that lead to the reorganization of conductive pathways through the material when a strain is applied (22). This reshaping of the percolating network leads to changes in the electrical resistance. Despite their potential for sensing technologies, using piezoresistive composites for biomedical sensors remains underexplored. To perform in biomedical devices and integrate into living tissue, piezoresistive sensors have to feature softness, sensitivity (i.e., High Gauge Factor), and monotonic response, particularly under dynamic conditions. Particularly in soft and bio-hybrid robots, they should display low piezoresistive hysteresis and viscous drift, while being suitable for tailorable fabrication processes (such as thermoplastic extrusion methods and fused-deposition modeling). The sensor materials should be biocompatible and have flexible and miniaturized designs that enable functional integration into biological environments. For example, sensors should compliantly interact with the target surfaces, soft tissue matrix, and structures of the biological culture environments if designed as wearable and implantable sensors, or integrated technologies for *in vitro* cell culture models. Such artificial sensory elements should not hinder tissue function and development. Finally, sensors should remain isolated from electrolytic components of the biological milieu.

To engineer a sensor that could meet all of these criteria, we selected piezoresistive composites based on carbon black (CB), which display lightweight, flexibility, high electrical conductivity, good mechanical properties, and safe manufacturing strategies. Piezoresistive composites based on CB-enriched styrene-ethylene-butylene-styrene (SEBS) thermoplastic elastomers can be manufactured into 3D structures via additive manufacturing and used as strain sensors, in which sensitivity, conductivity, and stretchability can be tuned by varying the properties of the thermoplastics (such as the shore hardness) and its formulation (such as the relative amount of the fillers) (24). Importantly, both CB and SEBS have been individually used for tissue engineering and other biomedical applications, demonstrating they can be safely incorporated with living materials (25–28). Nevertheless, combining CB and SEBS to form mechanosensors for optimized bio-integrated functionality has never been investigated. Our work demonstrated that these materials can combine with living tissue that is engineered *in vitro*, operate at a high sensitivity, and retain long-term functionality. We maximized the softness of our material and shaped it into a fiber design of small diameter (<0.25 mm) to enable versatile tissue-integrated designs and stabilize the bioelectronic interface between tissue and sensor. An insulation layer around the fiber was necessary to preserve the response signal from disturbance of other electrical inputs present in a cell stimulation setup. However, we showed that the insulation also protected the structural integrity of the sensor over time without affecting the ability of the tissue to interface with the sensor. Without the coating, sensor response characteristics, like the monotonic response, were diminished after prolonged exposure to the biological milieu. Our engineered tissue matrices and piezoresistive sensors can be assembled, without the engineered tissue losing its biological profile and the sensor decaying in its functionality.

The second important objective of our work was to create intelligent control of bio-actuators' motion. Current bio-hybrid systems lack system integration, which can be achieved by associating actuation with sensing and control strategy to generate operative autonomy. Such an embodied autonomy will enable performant, intelligent bio-hybrid robots. Understanding the bio-actuators' position and quantifying their movement are mostly performed via optical methods and dynamical analysis of recorded videos. For example, Iuliano et al. developed a tissue engineering platform that integrates optical fibers-based sensors to measure via light interferometry the displacement of a flexible cantilever induced by ring-shaped skeletal muscle tissue (29). If optical approaches are advantageous for multifunctional tissue engineering platforms, their implementation in robotic systems entails limitations. First, measurement inaccuracies are possible due to imaging limitations (*e.g.*, limited light transmission, as well as poor imaging resolution, video frame rate, and imaging processing). Second, using imaging-based measurements constrains the design of bio-hybrid robots, demands high-energy supply, and restricts the sensitivity to the detection of large deformation occurring in environments with controlled illumination. These systems require power-sourcing devices and precise positioning of the input and output components of the light sensor. Our piezoresistive sensor significantly simplifies the operative hardware in an energy-conservative system that does not need any additional power source, by which the sensor can connect to the setup required for cell electrical stimulation, and operates under any light condition. In addition, the tissue intimate contact allows one to detect not only motion but also tissue contractions expressed through minimal forces with a high accuracy and spatiotemporal resolution. By mentioning the need for feedback control mechanisms of bio-actuators, Kim et al. proposed to measure the contraction forces within a skeletal muscle-based bio-actuator with a mechanosensor made of liquid Galinstan embedded within soft elastomeric substrates (30). The cross-sectional and length deformations of the sensor that could be induced by the contracting muscle were expected to cause detectable resistance changes across the sensor. Even if the authors described the simulated performance of the sensor and theorized its applicability to a bio-actuator generating forces in a few mN ranges, the actual integration of the sensor and the tissue was not attempted, and no feedback control on the tissue functionality setup was reported. One of the main reasons could have been that the sensor design was composed of multiple discrete elements, including a liquid alloy component, an enveloping material, and inflexible copper wires. The presence of rigid components could cause strain shielding effects at the interface between tissue and electronics. With these premises, mounting it or co-developing it with a bio-actuator entails the risk of detachment or damage, with consequent release of the liquid conductive phase. Even if Galinstan is poorly cytotoxic at low dosages (31), its effects on cells at high dosages are yet to be investigated, implying that damaging the sensor could also compromise the bio-actuator's functionality. In contrast, our flexible sensor fiber based on piezoresistive elastomeric composite is consistent in its design, can easily integrate with both the engineered tissue and the reading interface, and can operate on a living muscle. It can be realized in designs that allow for optimal integration with the tissue, both before or following the tissue maturation process. Its constitutive materials (both in the core structure and at the surface) are highly biocompatible, thus limited tissue damage would be expected in the case of sensor damage. Finally, we prove our sensor's utility as a component for proprioceptive systems within soft matrices and conditions relevant to cell culture and cell electrical stimulation environment. Importantly, we generated a noise and/or artifacts-free mapping between the muscle contraction and relative resistivity change in the fiber that confirmed the detection of bio-actuators' deformations, which typically occur as micromotions. The map enabled the design of a gain closed-loop controller, which then realized the first, auto-determined behavior in a bio-hybrid actuator. In our closed-loop demonstration, we achieved that the switch-off actions of a controller could be effectively triggered by the tissue's contraction events with a high correspondence between the sensor signal peaks and the controller clearance outputs. Our data also indicates that the materials are safe to use with other cell types, including chemo-electroactive cells such as neurons. This observation expands the applicative horizons of this material to other uses in bio-mechanics and bioelectronics (such as multi-functional organ-on-a-chip devices), additionally benefiting from the sensor's elastomer-based composite nature, which combines a soft mechanical profile with the potential to cover a wide sensing range at low fabrication costs.

As further proof of the urgent need for solutions to mechanosensing in bio-systems, Zhao et al. recently reported a tissue-on-a-chip interface that enables intimate contact with mm-scale 3D skeletal muscle tissues for high-precision measurements of tissue contractility via a microscale strain sensor (32). Their sensor consisted of a planar system instrumented with strain sensors based on Cr/Au lines that connect to external control electronics. Despite being compliant and having the potential to be incorporated in electronic, optoelectronic, photonic, and microelectromechanical systems, this sensor only exhibits one-dimensional flexibility, prospectively limiting its implementation to well-developed planar devices. Interestingly, to amplify the resistance signal measured by the alloy sensor, the authors used a Wheatstone bridge, that provided highly accurate voltage readouts for minimal resistance variations. However, piezoresistive materials based on polymer composites can detect smaller forces than solid-state strain gauges (down to 1 mN and 40 mN for piezoresistive materials and strain gauges, respectively). Therefore, these materials could be used with a higher design freedom and provide an equivalent or more sensitive detection performance. In addition, the system's functionality was proven with optogenetically modified cells that respond to light stimulation, and not with wild-type muscle cells that require the application of electrical fields in the same space where the sensor is operating. Our insulated fiber-shaped sensor is functional in baths for cell electrical stimulation and offers the freedom to design tissue interfaces in 3D geometries inside soft matrices. These features open perspectives to realize complex sensor arrangements for implantable devices, 3D tissue models, and future motile bio-robots with portable designs.

Overall, our data represent a stepping stone for the generation of fully integrated biohybrid robots and a new starting point to research portable designs and onboarded multifunctionality in organic machines. We foresee that the next efforts will look at defining strategies for energy-efficient control systems, as well as the realization of more complex reaction schemes. Such schemes will be compatible with conditions and scenarios for real-life applications. For example, one next challenge will be the creation of adaptive control signals that increase the lifespan of bio-actuators' functionality by adjusting the stimulation paradigms to consider the natural deterioration or functional decay of the engineered tissue. Despite many advances in the field of bio-hybrid robotics, bio-hybrid systems are largely underdeveloped technologies and their performance cannot compare with traditional robotics under various metrics (5).

Our concept to sensorize bio-actuators will enable biohybrid robotics to overcome the challenge of full operative integration, inspire novel forms of biological machines capable of intelligent motion, and thus unlock the potential of biological robots, which, as endowed with autonomous operativity for real-life applications and will further express the unicity of their living component, through adaptability, self-healing, biodegradability, efficiency, and mechanical compliance. In a wider sense, the reported results can inspire technologies with direct application to pathophysiological 3D tissue models, micro-physiological systems, pharmaceutical research, and bio-hybrid system engineering.

### Extended Materials and Methods

#### Thermoplastic processing of the sensor fibers

Implant grade styrene-based tri-block co-polymer (TPS) was donated by Hexpol (Malmö, Sweden) in Shore hardness 20A, 40A and 50A. Carbon Black Ensaco 260G was obtained by Imerys (Paris, France). TPS and Carbon Black were mixed in 1:1 mass ratio, using the torque rheometer Polylab OS from Thermofisher (Karlsruhe, Germany). The temperature during mixing was 180°C. After the melt-mixing step, the composites were extruded in the form of fibers with a diameter of 0.2 mm, using the capillary rheometer RH7 from Netzsch (Selb, Germany). The temperature used during extrusion was 180°C. The diameter of the fibers was confirmed with the optical microscope Zeiss Stereo Discovery (Carl Zeiss Microscopy, Jena, Germany).

#### Dip coating of the sensor fibers

For the dip coating of the fibers, a liquid silicone rubber (LSR) EcoFlex 00-30 from Smooth-On (Macungie, PE, USA) was used in a 1:1 ratio. The fibers were coated by dipping them twice into the silicone rubber and leaving them to dry on aluminum foil. The dip coating process was repeated twice. The one end of the fibers was wrapped with Parafilm. The Parafilm was eventually removed to allow connection of the fiber with the Keithley 2450 electrodes with conductive copper tape. The resistance was measured alongside the length of the fiber to ensure that the coating process was successful.

#### Mechano-electrical characterization of the sensor fibers

The mechano-electrical testing was performed with the tensile testing machine Zwick Roell Z005 (ZwickRoell, Ulm, Germany). A 200 N load cell was used. The fibers were clamped with a pressure of 4 bar applied by pneumatic clamps. The electrical resistance was recorded during the tensile testing with a source meter (#2450, Keithley, Cleveland, USA). A voltage of 1 V and a sampling frequency of 10 Hz were used to characterize the fibers by tensile testing and 1000 Hz for the fibers integrated in the biorobots. A gauge length of 50 mm and a strain rate of 200 mm/min were used for the tensile test investigation. The relative resistance $R_{rel}$ was calculated by:

$$R_{rel} = \frac{R - R_0}{R_0}$$

where R is the measured resistance and $R_0$ is the measured resistance at the beginning of the tensile test.

The Gauge Factor (GF) was calculated with the following equation:

$$GF = \frac{\Delta R_{rel}}{\Delta \varepsilon}$$

Data were presented as mean values ± standard deviation with sample size ($n$ = 5). Statistical analysis was carried out using MS-Excel Software.

**Scanning electron microscopy**

The samples were fixed with 2.5% (v/v) glutaraldehyde in PBS solution at room temperature for 30 min and then rinsed three times with PBS. Then, the samples were dehydrated in an ascending series of ethanol solutions (30%, 50%, 70%, 90%, and 100% (v/v); 5 min in each solution), followed by three incubations of 10 min each in 100% ethanol, dried over a molecular sieve. After transfer into metal capsules, the samples were inserted into a critical point dryer (Tousimis 931) and the ethanol was substituted against liquid CO2. Then the samples were dried over the critical point of $CO_2$ (31 °C/73.8 bar). After the pressure was slowly released, the samples were taken out and mounted on SEM-stubs. For conductivity, the samples were sputter coated with 5 nm Pt/Pd (Safematic CCU-010). The examination was done in a JSM-7100F JEOL SEM at 3 kV by secondary electron detection.

**Functionality of the fiber embedded in soft matrices**

To understand the fiber functionality within soft tissues, we created various soft tissue phantoms and measured their shore hardness. To generate the phantoms, we cast various polymeric matrices into rectangular-shaped molds (size: 30×20×10 mm). Agar matrices were prepared by gelating different water solutions of agar (#A1296, Sigma Aldrich) at various w/v concentrations (from 0.5

to 2%). A matrix with no cells was prepared by mixing Matrigel basement membrane matrix (Corning, New York, USA) and Collagen (Cosmo Bio, USA) at a 1:1 ratio. The same matrix was tested immediately after myoblasts addition and after several days of tissue development, as explained in the section concerning skeletal muscle tissue fabrication. An analog Shore hardness tester (HB0 100-0, Sauter, Balingen, Germany) was used to measure the shore hardness at room temperature 2 hours after preparation. To test the functionality of the fiber within the matrix, the fiber was embedded in agar phantoms with shore hardness similar to the one reported for the matured tissue. Two embedding pre-tensioned designs were studied, with the fiber placed in straight or serpentine poses. Pre-tension on the fiber was created by fixing the fiber extremities in the straight pose and by using removable metallic clips to fix the loops. First, the fiber was inserted in the mold, and then polymers were cast and gelated around it. The surface adhesion and the flexibility of the fiber within stressed phantoms (1.5% w/v agar, size: 25×10×8 mm) under manual mechanical compression were shown with representative optical pictures.

**Cell culture**

The murine myoblast cell line C2C12 was obtained from the American Type Culture Collection (ATCC, Manassas, VA, USA) and was tested for mycoplasma (MycoAlert$^{TM}$ Mycoplasma Detection Kit, Lonza AG, Basel, Switzerland). Cells were cultured in monolayer at 37 °C in a 5% $CO_2$-containing humidified atmosphere in complete growth medium (GM), consisting of Dulbecco's modified Eagle's medium (DMEM, #D6429, Sigma–Aldrich) supplemented with 10% (v/v) heat-inactivated fetal bovine serum (FBS, #F7524, Sigma–Aldrich), 2 mM glutamine, 100 U $mL^{-1}$ penicillin, and 100 µg $mL^{-1}$ streptomycin (all from Thermo Fischer Scientific, Switzerland). Cells were grown at an initial seeding density of $5\times10^3$ cells/cm$^2$, and detached from flasks by trypsinization (Trypsin EDTA 0.25%, Sigma-Aldrich) at 70% of cell confluency. To induce cell myogenic differentiation, a differentiation cell culture medium was used, which consisted of DMEM containing 10% horse serum (Gibco), 1% penicillin-streptomycin, IGF-1 (50 ng/ml; Sigma-Aldrich), and ACA (1 mg/ml; Sigma-Aldrich). To prove the biocompatibility of the sensor components, other cell types were used. Murine fibroblasts (NIH 3T3 from ATCC) and motor neuron-like cells (NSC-34, TebuBio, Offenbach, Germany) were cultured in a high glucose formulation of DMEM supplemented with 10% FBS, as described above for C2C12. Human primary myoblasts (male donor, age 35) were obtained from Cook Myosite Inc. (Pittsburgh, PA, USA) and maintained using the MyoTonic Growth Media Kit (#MK-4444).

**Engineered skeletal muscle tissue fabrication and fiber integration**

To prepare cm-scale block-shape skeletal muscle tissue constructs, a polymer mixture composed of Matrigel basement membrane matrix (Corning, New York, USA) and Collagen (Cosmo Bio, USA) (ratio 1:1) was mixed with $1\times10^8$ cells/mL and manually cast in a rectangular block-shaped mold. The constructs were left in a cell incubator for 30 min, and then carefully unmolded, before being cultured in growth and differentiation medium for 3 and 6 days, respectively. To study tissue-fiber co-development, the fiber was inserted in the tissue construct during the biofabrication process by using a custom-made mold allowing for the correct positioning of the fiber (looped design, parallel to the main axis of the construct length). More details are shown in the Supplementary Materials. To fabricate ring-shaped skeletal muscle tissue constructs, cells suspended in a growth medium were mixed with Matrigel and fibrin (Sigma-Aldrich). The mixture was cast in a planar ring-shaped mold and cultured for growth and differentiation medium for 1 and 7 days, respectively, as described elsewhere.(33)

**Live/dead staining in 3D constructs**

To assess cell viability in the constructs, Live/Dead™ staining was performed immediately after bioprinting and at different time points during the culture time, following the manufacturer's instructions (Thermo Fisher Scientific; #R37601). Calcein and propidium iodide were excited at 488 and 561 nm laser wavelengths, respectively, and imaged on a confocal microscope (Zeiss LSM 780 Airyscan, Zeiss AxioObserver.Z1, ScopeM). Imaging analysis was performed on at least 3 images from each analyzed sample feature (≥ 3 samples/condition; ≥ 3 experimental replicates).

**Cell viability tests and cell growth in 3D Constructs**

Resazurin assays were used to evaluate the biocompatibility of the fiber materials and cell growth within the scaffolds. In biocompatibility tests, cells of different types were used. All cells were seeded in 96 well plates at an initial cell density of $5\times10^3$ cells/cm$^2$, and then incubated with different amounts of fiber materials (from 0.1 to 100 mg/ml) for variable durations (from 6 hours to 14 days). To perform the test, the culture medium was supplemented with 10% (v/v) of AlamarBlue reagent solution at 0.1 mg $mL^{-1}$ (#R7017, Sigma-Aldrich) and incubation was carried out for 2 h at 37 °C before the fluorescence signal ($\lambda_{Ex/Em}$ = 530/590) was measured with a Synergy H1 microplate reader (Biotek). Fluorescence intensity values were corrected for the background control (culture medium with resazurin). Cell viability was reported as a percentage index calculated against values calculated from control conditions lacking any added materials. Moreover, the Resazurin assay was performed to determine the metabolic activity of seeded cells in 3D constructs and confirm their proliferation. Briefly, the media was supplemented with the AlamarBlue reagent and after incubation (2 h, 37 °C), the media was collected and placed in a new well plate for reading fluorescence intensity values.

**Immuno-fluorescence in 3D constructs**

After *in vitro* culture, the constructs were fixed overnight in a 4% Paraformaldehyde solution at room temperature. After fixation, samples were rinsed in PBS (5 min, 3×), and stained for f-actin, MyoD, and Myosin Heavy Chain (MyHC). Briefly, samples were permeabilized with 0.1% Triton X-100 (20 minutes) and blocked with a 1% Bovine Serum Albumin (BSA) solution (1 hour). After extensive washing, the constructs were stained with the primary Anti-MyoD Antibody (#ZRB1452, Sigma-Aldrich, dilution 1:1000) and incubated for 4 hours with an anti-rabbit secondary antibody coupled with Rhodamine (#SAB3700846 Sigma-Aldrich, dilution 1:200). Cells were incubated in AlexaFluor 488 phalloidin (#R37110, Invitrogen) for one hour. 4′-6-diamidino-2-phenylindol (DAPI) was finally used to label the nuclei (dilution 1:1000; 15 minutes). All incubations were performed under gentle shaking and at room temperature. The staining of MyHC was performed by incubating the samples for 4 hours with the mouse anti-MyHC antibody (Myosin 4, eFluor™ 660, Clone: MF20, Affymetrix eBioscience™), which was used at a 1:10 dilution for a final concentration of 10 µg/mL. All images were taken using a confocal microscope (Zeiss LSM 780 Airyscan, Zeiss AxioObserver.Z1, ScopeM), using the same exposure time, and were analyzed in Fiji ImageJ. Differentiated myotubes in a specific microscopic field were observed under ×10 and ×20 magnification. Either the total number of nuclei or the number of nuclei within MyHC-positive myotubes was counted in 5 fields/sample. The fusion index was calculated as a ratio between the number of nuclei within $MyHC^+$ myotubes and the total number of nuclei. The images were randomly selected from at least 3 regions from each sample, and the experiments were replicated three times.

**Immunohistochemistry on tissue sections**

After *in vitro* culture, the constructs were fixed overnight in a 4% Paraformaldehyde solution at room temperature. Then, the samples were embedded in paraffin and sections of 4.5 μm thickness were cut with a microtome (Microm, HM430, Thermo Scientific). The sections were stained with Hematoxylin-Eosin (#GHS116 and #HT110116, respectively from Sigma Aldrich). The slides were mounted and imaged in Bright Field with a light microscope (Olympus CKX41, Olympus Schweiz AG).

**Secretome Analysis**

The amount of IL-4, IL-6, and Myostatin in the liquid media of constructs was quantified by collecting media aliquots at different time points and analyzing them via ELISA (incubation at room temperature for 2 h). Mouse IL-4 and IL-6 ELISA kits were purchased from R&D system (Minneapolis, MN, USA; #M4000B and #M6000B). Myostatin ELISA kit was purchased from LS Bio (Seattle, WA, USA; #LS-F35789).

**Sensing bio-actuator's contractions and motion analysis**

Six-well plates were coated with 2% agar (#A1296, Sigma Aldrich) to form a 0.5 cm-thick layer and the muscle tissue ring-shaped constructs were transferred into them with DMEM. The coated fiber was sterilized under UV radiation and then looped inside the muscle tissue rings. A pillar was inserted through the hole of the muscle ring into the agar bed in a vertical position. The fiber was anchored to the opposite side of the culture well and tended, resulting in a configuration of the muscle suspended between the tensioned fiber and the pillar. The extremities of the fiber were attached to a source meter (#2450, Keithley, Cleveland, USA) and electrodes were connected to a function generator (#DG812, Rigol, Beijing, China). A custom-made holder with two graphene electrodes was inserted into the culture medium, and electrical cell stimulation was performed by applying square pulses with 1Hz frequency at 4.5 V/cm (10 V; electrode distance: 2.2 cm) and 1% duty cycle. The experiment was recorded on a camera mounted on the microscope, and the videos were used to assess the motion performance of the biohybrid robots. Lateral displacements were calculated with Fiji ImageJ imaging analysis program.

#### Control system

To verify if the sensor system could be used in a closed loop controller, we analyzed the relative resistivity change measurements on the fiber obtained by applying square pulses with 1Hz frequency at 4.5 V/cm (10 V; electrode distance: 2.2 cm) and 1% duty cycle. Due to the high presence of noise on the obtained signal, three Gaussian $N(\mu, 1)$ filters were applied sequentially to the original signal. The filtered signal was then processed with a simplified thresholded Canny edge detector function to determine the presence of significant minima corresponding to the maximum contraction stages of the bio-hybrid setup. A threshold closed-loop controller was realized by feeding the data on an update loop as it became available and performing real-time processing on past measurements. The controller was configured to trigger a simulated signal at 1 Hz and analyze the rate of change of relative resistivity in the fiber to determine the shut-off point on each trigger cycle. The evaluation was performed on every measurement update (sampled at 16 Hz) and realized by applying a Gaussian $N(\mu, 1)$ filter on all the raw available measured data, followed by a weighted median filter (w = [1, 3, 5, 3, 1]), and an estimation of the rate of change relative to the fifth prior measurement and evaluated with a predefined threshold.

$$\frac{dR_{rel_t}}{dt} = R_{rel_t} - R_{rel_{t-5}}, t = 192, \ldots, 800$$

The threshold was set to -0.006 for both gathered samples to differentiate between true contractions and noise.

#### Data analysis and statistics

Data were analyzed with the Graph Pad Prism 9 Software. All variables are expressed as mean ± standard deviation (SD). Data were acquired from at least three independent experiments and three technical replicates unless otherwise stated. To assess statistically relevant differences between two experimental groups, the *t*-test was used ($P < 0.05$ and $P < 0.01$ are expressed as * and **, respectively). A general linear two-way ANOVA test was used to verify the hypothesis whether there were changes in various parameters over time among the experimental groups and to identify relevant variations among several experimental groups.

### Video Captions

**Movie S1. Controlled functionality of the bio-actuator stimulated in a closed-loop system.** Sensor fiber integrated into a ring-shaped bio-actuator with contraction response shaped by the activity of a controller set to trigger at 1 Hz. Contractile micromotions are mostly visible on the left side of the video, marked with an arrow.

**Movie S2. Static condition of the bio-actuator in the absence of applied electrical fields.** A sensor-integrated ring-shaped bio-actuator that does not move in the absence of electrical stimulation, thus not altering the controller behavior.

**Movie S3. Open loop of bio-actuators at 7 days of differentiation.** Sensor fiber integrated into a ring-shaped bio-actuator that responded to applied electrical pulses (20 Vpp; 1Hz) after 7 days of tissue maturation. Contractile micromotion is mostly visible on the left side of the field of view, marked with an arrow.
